# Improving Joint Estimation of Vital Rates in IPMs via Gaussian Processes and ABC

**DOI:** 10.1101/2025.07.02.662764

**Authors:** Zhixiao Zhu, Maria Christodoulou, David Steinsaltz

## Abstract

1. Developing population dynamic models that can flexibly adapt to different species remains a challenge due to the context-dependent nature of demographic studies. This work introduces ABC GP IPM, a novel Integral Projection Model (IPM) framework, aimed at mitigating key limitations associated with traditional IPMs, including (i) reliance on predefined vital rate models with restrictive assumptions, (ii) omission of potential vital rate interactions, and (iii) limited flexibility in incorporating domain- or user-specific knowledge. The purpose of this work is to enhance the adaptability and accuracy of IPMs across various ecological and evolutionary studies.
2. Our methodology integrates Gaussian Process (GP) models with Approximate Bayesian Computation (ABC). GP models introduce a non-parametric structure that accommodates a broader range of demographic patterns, reducing sensitivity to specific model choices. The incorporation of ABC methods allows IPMs to integrate additional population-level information, enabling potential interactions among vital rates to be reflected in model outcomes without requiring additional datasets. The information can be user-modified, facilitating the tailoring of IPMs to specific requirements or domain-specific knowledge. Recognising the inherent challenge of selecting appropriate summary statistics in ABC applications, we further propose a systematic procedure tailored to navigate this complexity effectively when applying ABC GP to IPMs.
3. The method demonstrated strong flexibility and adaptability across different modeling scenarios. In simulation studies, ABC GP IPM showed notable improvements over traditional GLM-based IPMs across multiple population metrics commonly used in biological research. When applied to real-world datasets, the method captured complex demographic patterns more effectively, supporting its potential as a practical and adaptable alternative to existing IPM frameworks.
4. The ABC GP IPM framework provides greater flexibility in model specification, accommodating potential non-linearities with fewer assumptions. It also facilitates the incorporation of user expertise and domain-specific knowledge, enhancing its applicability to context-dependent demographic studies. While offering a more adaptable modeling approach, the method remains accessible to users without specialized expertise in statistical modeling. Importantly, ABC GP IPM delivers these improved outcomes using identical datasets as standard IPMs, facilitating easier adoption for studies that have applied traditional IPMs.

## 1 Introduction

Life cycles define an organism’s journey from birth to death, highlighting key transitions like from juvenility to adulthood and from non-breeding to breeding. These stages are quantified mathematically by vital rates, such as survival and growth rates. Influenced by environmental conditions and inherent traits, the rates continuously interact, illustrating the biological trade-offs (Stearns (1989); McNamara & Houston (1996); Obeso (2002)). These demographic processes and trade-offs sculpt unique life paths for individuals. Over time, these paths converge, forming cohorts and populations, shaping the evolutionary history of the species.

Integral Projection Models (IPMs) serve as a fundamental tool in these demographic studies, offering key insights into natural selection and improving predictions of population trends (Picó et al. (2008)). Demographic studies are inherently context-dependent, with variations across species and research objectives. To cater to these diverse requirements, IPMs have developed to incorporate, for example, more detailed multi-dimensional state variables (Childs et al. (2003); Coulson et al. (2010); Stubberud et al. (2019), factors like population density (Rose et al. (2005); Rebarber et al. (2012); Coles et al. (2023)) and environmental stochasticity (Childs et al. (2004); Dahlgren et al. (2016); Félix-Burruel et al. (2024)). These developments reflect the ongoing efforts to tailor IPMs to the complex realities of ecological systems. Nevertheless, current IPMs still face challenges that compromise their ability to produce reliable population projections. There are three key limitations that we wish to highlight, and that we seek to address in this work.

### (1) Reliance on Predefined Vital Rate Models with Restrictive Assumptions

Traditional IPMs typically employ (generalised) linear and mixed-effects models to estimate vital rates (Easterling et al. (2000b); Ellner & Rees (2007)), favoured for their mathematical tractability and straightforward interpretability. While Bayesian extensions have been developed to better address uncertainties (Rees & Ellner (2009); Merow et al. (2014b); Messerman et al. (2023)), both parametric schools — Bayesian and Frequentist — tend to oversimplify or misrepresent the relationships between vital rates and state variables as linear or fixed, despite substantial evidence suggesting otherwise in nature (Lingjærde et al. (2001); Martorell & Martínez-Ballesté (2019); Bonnaffé & Coulson (2022)). Non-parametric approaches like splines (Dahlgren et al. (2011); Ellner et al. (2016); Macdonald et al. (2023)) provide an alternative when linearity is inappropriate, but their lack of probabilistic outputs limits their effectiveness. This limitation shifts the focus from a broader distribution that encapsulates inherent variability of vital rates to a narrow, precise, but potentially misleading point estimate (Ellison (2004); Hobbs & Hilborn (2006); Elderd & Miller (2016)).

### (2) Omission of Potential Vital Rate Interactions

Traditional IPM strategies often model vital rates separately before integration into the comprehensive IPM, assuming that accurately modeling each rate will result in a reliable overall population model. Such a piecemeal approach stands in contrast with the reality that vital rates are often interdependent, like reproductive costs and maturity thresholds (Kelly (1992); Hilborn et al. (2006); Struckman et al. (2019)). While some studies have employed Bayesian frameworks to detect such interactions through the joint distributions of multiple vital rates (Elderd & Miller (2016)), the likelihoods for the vital rates typically share no parameters, making their joint optimization equivalent to maximizing each separately (Plard et al. (2019)).

Interactions among vital rates can occur on different scales, ranging from interactions within individuals due to biological heterogeneity to broader patterns that emerge at the population-level (Vindenes et al. (2008)). Since IPMs are designed to describe population dynamics, it is important to account for interactions that arise at the population scale. However, such interactions, which shape broader demographic patterns, may not be well captured when vital rates are estimated separately using individual-level data.

To gain a more comprehensive understanding, population-level insights should also be incorporated, reflecting multiple vital rates and their potential interactions. For example, grouping individuals based on their status — such as growth or shrinkage among breeders and non-breeders — can help capture aggregated effects of vital rates across different groups. Incorporating such information provides additional perspectives on how vital rates collectively influence population dynamics — offering insights that traditional, independent analyses in IPM may overlook.

### (3) Challenges in Incorporating Domain Knowledge

Lastly, present-day IPM structures offer no obvious entry point to integrate users’ empirical insights, and to adapt to new research questions. Researchers accumulate valuable background knowledge about their study species through years of dedicated investigation. Incorporating their expertise could contribute to more realistic and well-informed population models. However, translating detailed biological knowledge into statistical frameworks remains challenging, due to the nuance of scientific insight, complexities of model formulation, coding, and the practical implementation of sophisticated, context-specific models.

We propose a novel methodology that enhances the robustness of the IPMs. Our method begins by replacing traditional base models with a Bayesian non-parametric model: the Gaussian process (GP) model. This model introduces a new level of flexibility, capturing complex dependencies without predefined functional forms and providing probabilistic outputs to better reflect natural variability (Williams & Rasmussen (2006)). We rely upon Approximate Bayesian Computation (ABC) methods, which enable researchers to infuse specific knowledge about the system being analyzed during model synthesis. The ABC reduces the complexity of translating intricate biological insights into statistical models, making the method more accessible to non-experts in modeling while allowing customization of IPMs to specific needs. As long as models can be represented by a data-generating stochastic process, researchers can turn them into applicable IPMs. Our approach utilises summary statistics in ABC that incorporate population-level data, helping capture vital rates’ joint behaviours at the population scale.

There are existing methods, in particular the hierarchical modeling framework for IPMs recently introduced by Fung et al. (2022), that allow for interdependencies among vital rates. However, this approach still relies on parametric assumptions, making the results sensitive to the chosen model structures. Integration of population-level data is also possible with the method proposed by Plard et al. (2019). This method relies on parametric assumptions to link individual- and population-level data. It simplifies implementation, compared with ABC, but at the cost of potentially introducing biases from the initial model assumptions, which are much less prescriptive in the data-driven ABC approach.

We begin by describing, in Section 2 the foundational methods underlying our study, including IPMs, GP, and ABC. We then present the enhanced framework that we call ABC GP IPM, highlighting the complementary roles of ABC and GP in addressing key limitations of traditional IPMs. Recognizing the challenge of selecting appropriate summary statistics in ABC applications, we propose a systematic procedure in Section 3 to navigate this complexity effectively. To evaluate our approach, Section 4.1 compares traditional IPMs with ABC GP IPM using simulated datasets with known “truth”, employing population metrics grounded on the *stable population growth theory*. This validation step is crucial for assessing the robustness and reliability of our method under controlled conditions. Building on these insights, Section 4.2 extends our methodology to real-world datasets of *Cryptantha flava*, demonstrating its practical viability in forecasting population trends. Finally, the report concludes with insights on the interpretation of our proposed method (Section 6).

## 2 Methods

### 2.1 The study species and the dataset

We focus on *Cryptantha flava*, sampled near the Red Fleet State Park in northeastern Utah, USA (Lucas et al. (2008); Salguero-Gomez et al. (2012)). *C. flava* was selected due to its extensive coverage in prior research with well-documented life history traits (e.g., Casper (1988); Lucas et al. (2008); Salguero-Gomez et al. (2012); González et al. (2016); Evers et al. (2021)), providing a solid foundation for our simulations and real data analyses. The dataset is comprehensive, spanning a considerable temporal range from 2003 to 2012, and includes detailed tracking of over 1,000 individuals. These attributes make *C. flava* an ideal candidate for testing and comparing the robustness and flexibility of our IPM methodology.

The plant adjusts its size (measured by the total number of rosettes) through either growth, by producing new rosettes from axillary meristems, or shrinkage, by losing existing rosettes from flowering or mortality. This non-independent relationship underscores the necessity of distinct growth models for reproducers and non-reproducers. Additionally, to better model population dynamics in interaction with the natural system in the real case study, we integrated environmental factors from a nearby weather station (USW00094030, NOAA (2013)). Specifically, we incorporated three annual weather variables to summarise inter-annual environmental variation: the mean daily maximum temperature, the mean daily minimum temperature, and the mean daily precipitation, each averaged across the year (Easterling et al. (2000a); Zou et al. (2009); Huxman et al. (2004)). These variables were included as covariates in our vital rate models, following common practice in IPMs involving environmental stochasticity (Dahlgren et al. (2016); Ellner et al. (2016); Félix-Burruel et al. (2024)).

### 2.2 Integral projection models

IPMs assume that individuals in the target population can be completely summarized by an attribute *z*, which can be multidimensional with both discrete and continuous variables (such as age-size) (Rees et al. (2006); Ellner & Rees (2006)). In IPMs, with current attribute *z*, a population at the next time step with attribute *z*′ can be described by

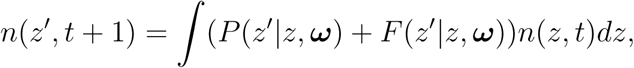

where ***ω*** represents environmental factors affecting the *survival-growth kernel*, *P* (*z*′|*z,* ***ω***), and the *reproduction kernel*, *F* (*z*′|*z,* ***ω***).

Tailored to *C. flava*’s life cycle, *P* and *F* can be further separated into individual vital rates to align with the actual life cycle events. *F* (*z*′|*z,* ***ω***) starts with considering the probability of breeding, *p_f_* (*z*|***ω***); conditional on flowering, a state *z* individual would produce an average of *n_f_* (*z*|***ω***) flowering stalks; each stalk leads to a certain count of seeds establishing and growing to size *z*′ by the next census, *r_est_r_size_*(*z*′). That is, *F* (*z*′|*z,* ***ω***) = *p_f_* (*z*|***ω***)*n_f_* (*z*|***ω***)*r_est_*_|_***_ω_****r_size_*(*z*′ ***ω***). The survival-growth kernel, *P* (*z*′|*z,* ***ω***), considers the reproduction’s influence on growth dynamics. It assumes that an individual with current state *z* will survive to the next year with probability *s*(*z*|***ω***), but reproducing or not would lead it to different fates in growth, *G_f_* (*z*′|*z,* ***ω***) or *G_nf_* (*z*′|*z,* ***ω***). That is, *P* (*z*′|*z,* ***ω***) = *s*(*z*|***ω***)*p_f_* (*z*|***ω***)*G_f_* (*z*′|*z,* ***ω***) +*s*(*z*|***ω***)(1 − *p_f_* (*z*|***ω***))*G_nf_* (*z*′|*z,* ***ω***). The sequence of life cycle events in these proposed sub-kernels, *F* (*z*′|*z,* ***ω***) and *P* (*z*′|*z,* ***ω***), align with the actual census process conducted in Salguero-Gomez et al. (2012). We employ these IPM sub-kernels in both future simulations and real dataset case studies to better capture observed demographic processes.

### 2.3 Vital rate in IPMs and Gaussian Process models

In IPMs, vital rate functions serve as the critical link between individual-level observations and population-level projections. These functions capture and summarize variations and responses among individual observations, which are then incorporated into sub-kernels *P* and *F* to facilitate population projections. As highlighted by Merow et al. (2014a), “IPM predictions can be only as good as the assumptions and inferred transitions of these vital rate models”. The vital rates thereby need to be fitted with caution to avoid misleading population-level inferences.

Traditionally, vital rates in IPMs have been parameterized using conventional statistical methods, like linear regressions for growth and logistic regressions for flowering probability etc.. These parametric methods assume that the data are generated from distributions that follow tightly constrained functional forms, and then determine the optimal parameters within these constraints. While effective in some scenarios, these methods can be restrictive and misleading when the true relationships in the data deviate from the pre-assumed models. Modern computational tools allow models to learn functional relationships directly from the data, minimizing the need for predefined functional forms. One such tool is the GP, also known as *Kriging method* in geostatistics. GP models are well-established and provide interpretable frameworks (Williams & Rasmussen (2006)). Although GPs have been increasingly applied in ecological studies, like species distribution modeling (Golding & Purse (2016); Ingram et al. (2020)), to the best of our knowledge, their integration into IPMs remains unexplored. Dahlgren et al. (2011) have demonstrated that cubic splines perform well for IPM estimation when the assumption of linear relationships between vital rates and state variables is not satisfied. GP models, as a generalisation of splines (see Kimeldorf & Wahba (1970); Bay et al. (2016)), are a natural extension for capturing complex nonlinear relationships in IPMs.

A GP is defined as an infinite collection of random variables, where any finite subset follows a multivariate normal distribution (MVN). In practice, we typically evaluate a GP at a finite set of input points, yielding a finite realization. Given a hyper-parameter *θ* ∈ P, such a realization *f_GP_* can be written as:

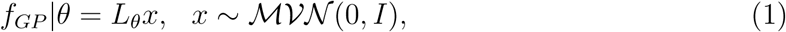

where *I* is an identity matrix and *L_θ_* is the Cholesky decomposition of the GP kernel matrix 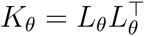.

Although each realization of a GP gives function values at a finite set of inputs, the GP defines a distribution for any such set. These distributions are not isolated: they are constructed using the same GP kernel and agree on overlapping inputs.

This structure allows us to interpret the GP as a distribution over functions, rather than just over values at specific input locations. It, thereby, highlights the role of GPs as Bayesian non-parametric models, providing a flexible distribution over numerous potential functions that align with observed data, rather than being restricted to a specific distributional form. The kernel matrix *K_θ_* represents the covariance structure of the GP. The strength of GPs lies in their ability to measure data point similarities through the kernel *K_θ_*. Generally speaking, GP models assume that data points closer together in the input space are more similar in the output space. The input space similarities are algebraically transformed to reflect output space correlations, enabling GPs to provide a predictive distribution for new data points based on observed datasets. To mimic such complex ‘correlations’ observed in real data, GP models potentially have an infinite number of parameters, allowing the size of the models to grow with the data size (Li et al. (2019)). Although GPs are fundamentally non-parametric, they incorporate hyper-parameters *θ* to regulate model complexity, thereby preventing over-fitting.

### 2.4 Approximate Bayesian Computational Methods

ABC is a set of sampling methods for Bayesian inference that has been effectively applied across various ecological and biological studies (e.g. Jabot & Chave (2011); Scranton et al. (2014); Daly et al. (2017); Zhang et al. (2017); Ruiz-Suarez et al. (2020); Lux (2021)). Traditional Bayesian inference requires explicit calculation of the likelihood function to provide an exact posterior distribution. However, in many biological studies, while it is feasible to simulate datasets, converting these simulations into an evaluable likelihood function is often difficult or impractical. In these scenarios, ABC is particularly useful for facilitating Bayesian analysis. It bypasses the need for a known likelihood function by using simulations to estimate the likelihood. The core process of ABC involves four main steps:

1. **Simulate Data:** Sample parameter values from a prior distribution, *π*(***θ***), and generate simulated datasets.
2. **Compare to Observed Data:** Compare each simulated dataset to the observed data. Typically, this comparison employs a set of summary statistics instead of the full dataset to reduce dimensionality and computational demands.
3. **Accept or Reject:** Accept a parameter sample if the simulated summary statistics, ***s***^∗^, are closely match those observed summary statistics, ***s****^obs^*, based on predefined thresholds; otherwise, reject.
4. **Posterior Approximation:** Treat the accepted parameter values as samples from an approximation of the posterior distribution, *π*(***θ***|***s****^obs^*) ≈ *π_ABC_*(***θ***|***s****^obs^*).

ABC’s effectiveness hinges on the choice of summary statistics and the distance threshold, both of which significantly influence the quality of the posterior approximation. If the summary statistic ***s*** for the parameter ***θ*** is *(Bayes) sufficient* (Cox & Hinkley (1979)) and the simulated and observed summary statistics match perfectly, then ABC no longer provides a mere approximation. Instead, it recovers the exact posterior distribution, i.e. *π_ABC_*(***θ***|***s****^obs^*) ≡ *π_ABC_*(***θ*** *y_obs_*) *π*(***θ*** *y_obs_*). Section 3 outlines a proposed workflow for selecting ABC summary statistics systematically.

Traditional ABC methods, from simple rejection-based approaches (Tavaŕe et al. (1997)) to more advanced techniques like ABC-MCMC (Marjoram et al. (2003); Wegmann et al. (2009); Lenormand et al. (2013); Andrieu et al. (2018)) and ABC-SMC or ABC-PMC (Beaumont et al. (2009); Toni et al. (2009); Sisson et al. (2009)), have greatly improved parameter inference for complex models. However, these methods often struggle with complex combined models, like IPMs — large frameworks composed of multiple sub-models. The “curse of dimensionality” becomes a major challenge, particularly when identifying suitable global perturbation kernels for each sub-model of vital rates. Poorly chosen kernels can lead to excessive rejection rates, substantially increasing computational demands.

To ameliorate the limitations of traditional ABC methods in complex scenarios, a modified ABC-PMC (Population Monte Carlo) approach has been proposed (Zhu et al. (2025)). This method incorporates pre-calculated MCMC samples from each sub-model to refine the ABC prior distribution and perturbation kernel. By leveraging insights from simpler model components and employing an adaptive weighting strategy that consolidates summary statistics into a single metric (Prangle (2017)), this approach enhances computational efficiency and facilitates parameter inference across multi-component models.

Here we have applied this modified ABC-PMC methodology to IPM within the ABC GP framework. This ABC-PMC method, detailed in Algorithm 1 and Table 4 in Appendix A, operates as follows:

1. **Initialisation:** Generate an initial set of particles from an informative prior based on pre-calculated MCMC samples. Simulate datasets and compute initial distances using summary statistics, assigning equal weights to all particles.
2. **Iterative Process:**

- For each step *c* = 1 to *C*, resample particles based on their weights from step *c* − 1 and perturb them by randomly altering one vital rate parameter using MCMC samples.
- Simulate a new dataset from each perturbed particle and recalculate the distance to the observed data.
- Accept particles if their recalculated distance is below a dynamically computed quantile-based threshold. Update and normalise weights of accepted particles.
3. **Output:** Repeat until the final iteration, producing a weighted set of particles that approximate the posterior distribution.

### 2.5 Individual-based models

ABC methods rely on simulations. In this study, we generated all simulated population datasets using Individual-based models (IBM) (see reviews in Grimm et al. (2006); DeAngelis & Grimm (2014)). The IBM is a population simulation technique that tracks the life trajectory of each individual and takes demographic stochasticity into account to mimic real life. In our IBM implementation, demographic events occur in accordance with the life cycle and are treated as random based on probabilities parameterized from the same GP/GLM modelled vital rates. These ensure the IBM is realistic in the sense that the simulated demography is roughly comparable to that of the observed population (Ellner et al. (2016)). Our use of IBM in ABC generates population outcomes under specific hyper-parameter values of IPMs, making them ready for incorporation into the sampling framework of the ABC-PMC algorithm

### 2.6 Enhanced IPMs

In response to the three identified limitations, we propose to enhance IPMs by integrating GP and ABC. Traditional Bayesian IPM fitting typically involves independently fitting GLMs to vital rates; these GLMs are then combined to assemble IPMs by drawing random samples from MCMC outputs (grey arrows in Figure 1). Our approach, illustrated by the blue and orange arrows, modifies and extends this traditional IPM construction process.

1. **Integration of GP models:** We begin by replacing traditional base models, such as GLMs, with GP models. This approach (indicated by the blue arrows) follows the same structure as traditional IPM practices, modeling each vital rate independently using individual-level data under a Bayesian framework, with the only difference being the use of GPs instead of GLMs.
2. **Constructing IPMs with ABC-PMC:** Instead of assembling IPMs by drawing independent MCMC samples for each vital rate, we employ the ABC-PMC sampler to construct IPMs (orange arrows in Figure 1). This approach retains only the combinations of vital rates that are able to ‘reproduce’ the observed characteristics.

**Figure 1:**
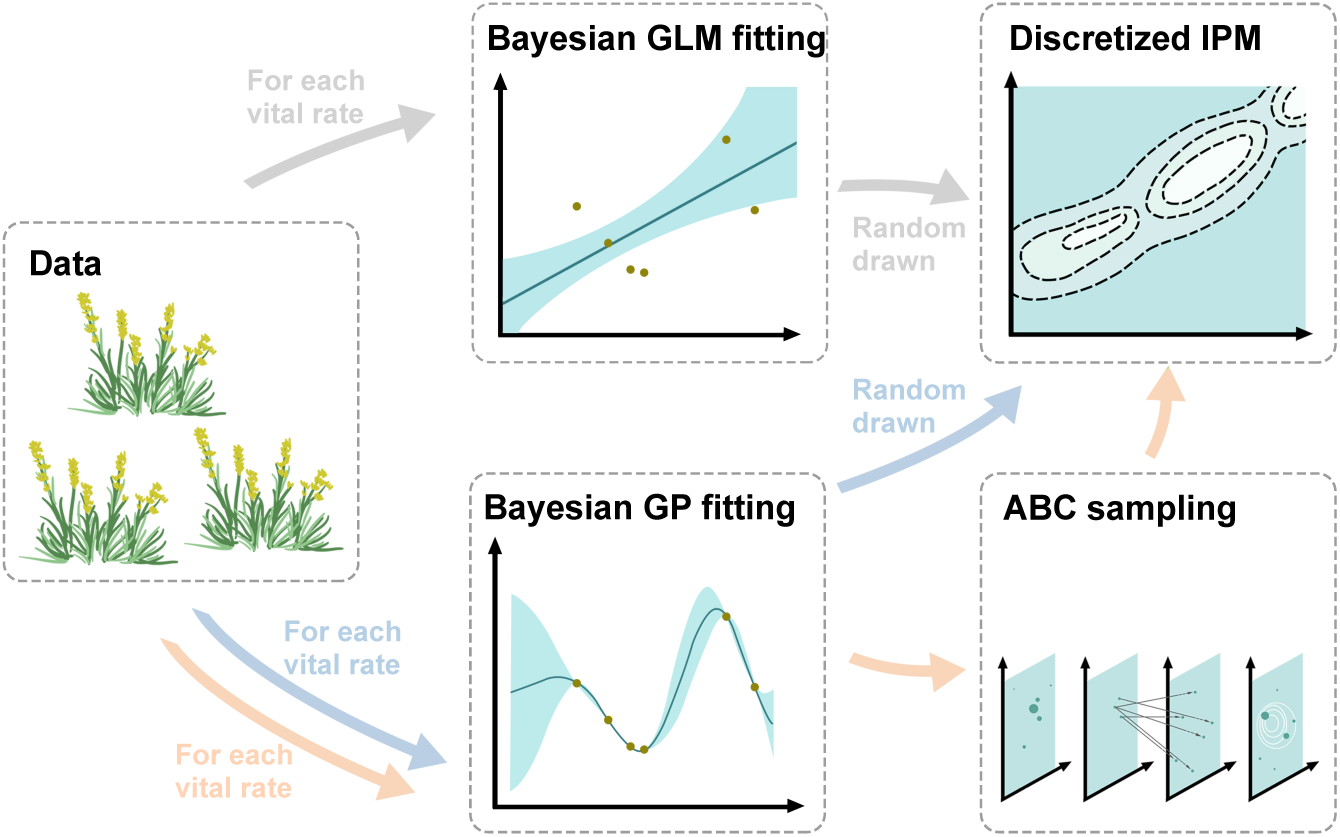
Workflow chart illustrating the enhanced IPM construction Process: This diagram demonstrates the integration of GP models and ABC in the development of IPMs. Grey arrows depict the steps involved in constructing a traditional Bayesian IPM. Blue arrows highlight the replacement of standard GLMs with GP models. Orange arrows illustrate the innovative application of the ABC GP methodology to synthesise IPMs more cohesively. Please note that all data points and content in this chart are for illustrative purposes only and do not represent any actual data values or practical implementations.

The incorporation of GP models introduces flexibility and provides probabilistic outputs, handling inherent nonlinearities in natural systems. This setting mitigate structural sensitivity issues (Wood (2001)) and is particularly beneficial when data is limited. Additionally, the integration of ABC helps us to incorporate population-level data, capturing the joint behaviours of vital rates essential for population dynamics.

## 3 A systematic workflow for selecting summary statistics

Identifying *(Bayes) sufficient* summary statistics for *y_obs_* is challenging in real-world applications. ABC methods thereby, in typical practice, settle for the next best thing, assuming that *π*(***θ***|*y_obs_*) can be reasonably approximated by *π_ABC_*(***θ|s****^obs^*) when only *insufficient* summary statistics are available. In biology, interpretable summary statistics are preferred, for reflecting users’ practical understanding of the studied system (Newman et al. (2022)). In areas with extensive ABC applications, summary statistics can be selected from previous studies. However, in new applications of ABC, selecting the appropriate summary statistics requires more careful consideration. For example, the fixation index could be very informative in the context of one model but less so when switching to a different model (Wright (1949); Nielsen & Wakeley (2001)).

An ABC algorithm requires that essential features of the data be encapsulated in summary statistics, and that distance metrics be defined to appropriately quantify dissimilarities between datasets based on these summaries. Different metrics can lead to varied acceptance results, impacting the quality of posterior approximations (Bernton et al. (2019)). Identifying the ideal metric often involves experimenting with multiple options (Li & Jakobsson (2012)). To address these challenges in our novel ABC application in IPMs, we propose a set of candidate statistics informed by the system’s background, as detailed in Table 1. For the same row of Table 1, each item in the left column can be paired with an item on the right to create a unique combination of summary statistic and distance metric, producing 31 combinations in total. For brevity, we also refer to these combinations as “summary statistics” throughout the rest of the paper.

**Table 1:**
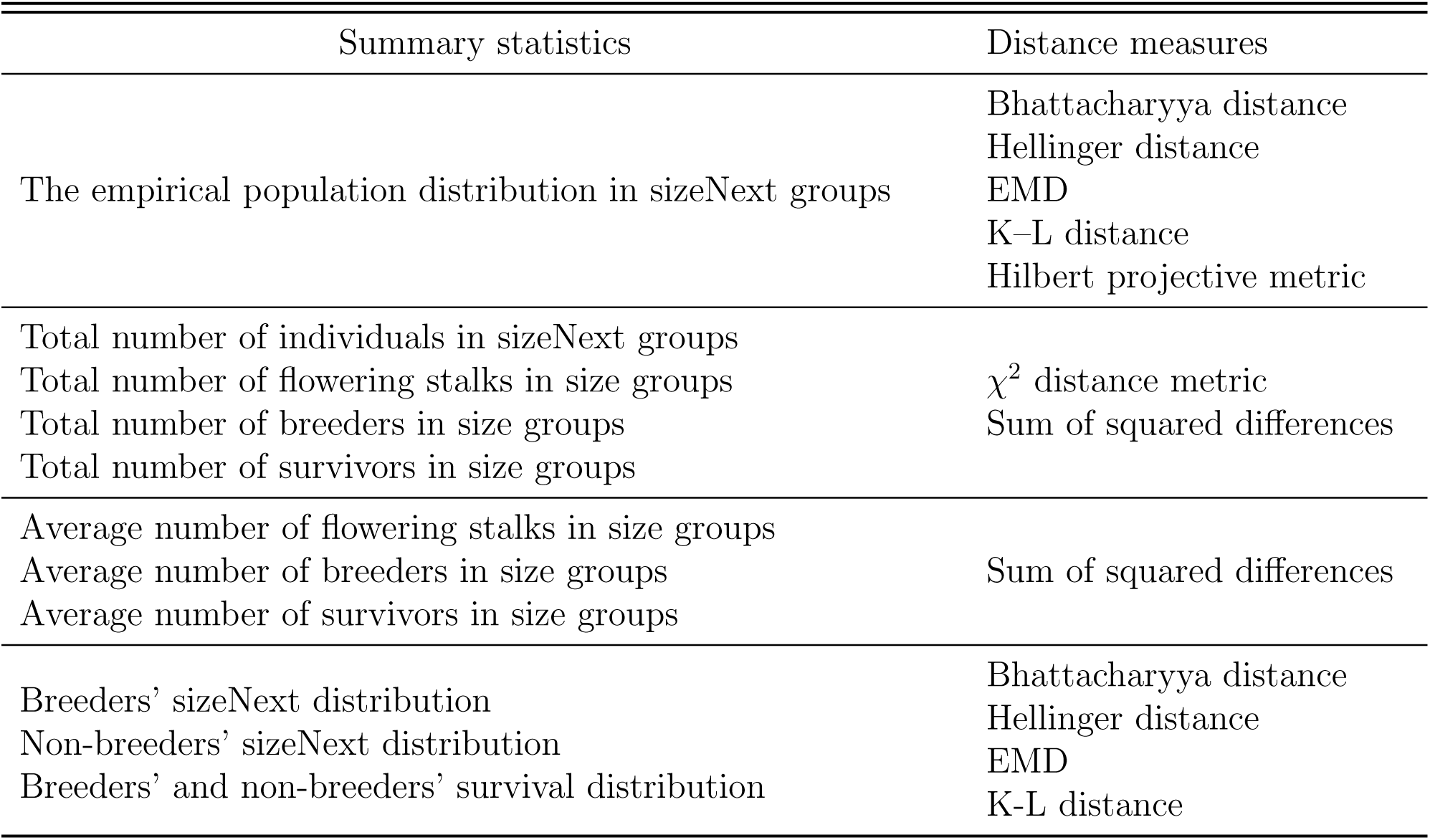
The list of candidate summary statistics and the corresponding distance measures in our study. To calculate the majority of candidate summary statistics under consideration, individuals need to be grouped first according to their size. sizeNext represent the individuals’ size in the next year. Please see Appendix B.1 for a full description.

To determine the optimal subset of summary statistics, we follow Scranton et al. (2014) to evaluate how well each candidate distinguishes dataset pairs generated from models with ‘slightly different’ hyperparameters. We use the Maximum Likelihood Estimates (MLEs) from the fitted model as a baseline and simulate datasets based on these estimates. Perturbed versions of the MLE model, generated by adjusting hyperparameters, are then used to simulate corresponding datasets for the comparisons.

The standard perturbation approach would vary one hyperparameter at a time, while keeping others fixed. This approach runs into difficulties when applied to GPs, in particular:

- **Hyperparameter interaction:** Hyperparameters in GPs often interact in complex, non-linear ways, where small changes in one can affect others and may amplify their overall impact on the model. Standard perturbation methods fail to capture these interactions, resulting in a mismatch between the scale of the perturbation and the model’s response.
- **Dependency on hyperparameter magnitude:** The impact of perturbations depends heavily on the initial hyperparameter value. Different regions of the hyperparameter space may exhibit varying levels of sensitivity, making it difficult to generate consistent and reasonable perturbations.

See Appendix B.2 for an illustrative example, and an empirical experiment in Appendix C.2 showing that perturbations of similar magnitude can result in highly varied model fits.

Given these challenges, we adopt an alternative approach, using Negative Log Posterior Probability (NLPP) scores to define model similarity. In the cases of flat prior distributions, since NLPP scores are derived from the log-likelihood of the training data points, similar NLPP scores indicate that the training data points are similarly probable under each model. When users have additional beliefs or knowledge about the hyper-parameters’ values before observing the data (informative priors), the NLPP integrates this belief, reflecting it in the score. By keeping the training data points fixed, this method makes stable and consistent comparisons in GP models, alleviating the limitations of parameter-driven perturbation approaches.

Under this setting, models with comparable NLPP scores are considered similar, regardless of differences in their hyperparameter values or predictive surfaces. This method involves varying multiple hyper-parameters simultaneously to create new models with NLPP scores close to the original.

We then evaluate each candidate summary statistic based on its ability to distinguish datasets generated from the model with the lowest NLPP from its perturbed variants. Each summary statistic acts as a classification tool, with its effectiveness assessed through the area under the receiver operating characteristic (ROC) curve (AUC). The AUC measures classification performance by comparing true positive and false positive rates across varying thresholds. An AUC of 1.0 indicates perfect classification, while an AUC of 0.5 corresponds to random guessing (Hastie et al. (2009)). The detailed workflow for this approach is outlined in Workflow 1. Notice that, to illustrate the procedure, we describe the workflow using specific numbers (e.g., 1,000 simulations, 2% closest samples). These are illustrative and can be adjusted depending on computational resources and application needs.

##### Workflow 1

For each vital rate, we repeat the following steps while keeping the other vital rate models fixed at their MLEs:

1. **GP Model-fitting:** Fit a GP model to the observed dataset within a Bayesian framework. This yields, e.g., 5,000 sets of MCMC samples for hyper-parameters for the target vital rates.
2. **Optimal NLPP:** Determine the NLPP score for each MCMC sample. The one with the smallest score is denoted as ***θ****_opt_*.
3. **Simulations at the Optimum:** With ***θ****_opt_*, simulate, for example, 1,000 pairs of complete datasets through one-step forward projection independently (i.e., 2,000 datasets). For each pair, calculate distances using all 31 candidate summary statistics (listed in Table 1), yielding 31 distributions 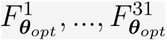 that represent baseline variability under the optimum.
4. **NLPP Perturbations** & **Re-simulations:** From the MCMC samples, select the top 2% closest to ***θ****_opt_* in NLPP (e.g., 100 samples), denoted as {***θ****_k_*}*_k_*_=1_*_,…,_*_100_. Treat them as NLPP perturbation outcomes for ***θ****_opt_*. For each ***θ****_k_*:

- Simulate 1,000 datasets under ***θ****_opt_* and another 1,000 datasets under the ***θ****_k_*.
- Pair corresponding datasets (by index) and compute distances across the summary statistics (as in Step 3). For each statistic *i* and each perturbation *k*, this yields a distribution of distances 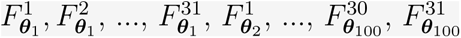.
5. **Summary statistics Selection:** For each statistic *i*, compute the AUC value between 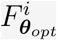 and each of 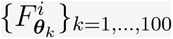, denoted as 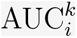. Then count how many times 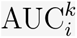 exceeds a specified threshold *ρ* across all *k*, 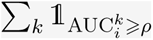. The statistic with the highest count is selected as the most sensitive for the corresponding vital rate.

As illustrated in step 5 in Workflow 1, we prioritize summary statistics that demonstrate consistent performance across different hyperparameter settings (through 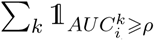), rather than those that perform well only in specific situations. This is because, during ABC sampling, model parameters vary across iterations, and statistics that remain informative across a range of plausible models are expected to perform more reliably. Depending on specific modeling goals, alternative selection criteria could also be considered.

In step 5, we set a classification threshold on the AUC and assess how frequently each summary statistic exceeds it across perturbed models. A threshold around 0.75 is often used as a reference for acceptable classification performance (e.g., Hosmer Jr et al. (2013)), although, in practice, different thresholds may be applied depending on the study objectives (e.g., Iyer et al. (2016); Gail & Pfeiffer (2018); Ç orbacıoğlu & Aksel (2023)). Summary statistics that surpass this threshold more frequently are considered more robust and are selected for our subsequent ABC sampling.

## 4 Case Studies

### 4.1 The simulation study

To demonstrate the application of ABC GP in IPM, we conducted a case study using simulated datasets for *C. flava*. Taking advantage of the known truth in simulated data, we evaluated how a misspecified linear model performs when the true relationships are nonlinear or, conversely, how model-fit accuracy suffers when the more robust procedure is applied that ignores the “true” linear relationships. We simulated two datasets under linear and nonlinear assumptions and fitted each using the strategies shown in Figure 1. These strategies include:

1. **GLM:** fitting vital rate through Bayesian GLMs; IPMs are constructed by randomly assembling one posterior draw per vital rate from their respective MCMC samples;
2. **GP:** fitting vital rate through Bayesian GPs; IPMs are constructed by randomly assembling one posterior draw per vital rate from their respective MCMC samples;
3. **ABC GP:** fitting vital rate through Bayesian GPs; IPMs are constructed through ABC-PMC.

We then analysed the resulting IPMs, facilitating for a comprehensive comparison of the three modelling strategies.

#### 4.1.1 Simulation Setup and Model Fitting

We initiated our experiment by applying GP models and GLMs within a frequentist framework to fit the real dataset (aggregated from the three years from 2008 to 2011). Using these fitted models (Table 2), denoted *M*_gp_ and *M*_glm_, we conducted IBM simulations to project population dynamics over a decade, creating two simulated datasets: *D*_gp_ and *D*_glm_. These datasets were treated as surrogate ‘truths’ for the subsequent simulation case study.

**Table 2:** Estimated demographic functions in GLMs and GP models for *C. Flava*. Here, *z* represents the log-transformed size; *f*_gp_ follows GP with Squared Exponential Kernel measuring input differences, 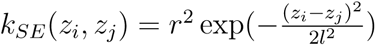 and zero mean functions. As information about mother plants is frequently missing, offspring size *r_size_* and establishment probability *p_est_* were taken to be independent of maternal size, for both the GLM and GP methods. We assumed *r_size_* and *p_est_* were known and fixed for all fitted IPMs to maintain a consistent baseline across models. A Gamma distribution (*α* = 0.706*, β* = 1.329) was used, replacing the traditional Normal distribution, to fit the size of new recruits, *r_size_*, due to the absence of negative sizes. The probability of offspring establishment (*p_est_* = 0.210) was calculated by dividing the number of newborns by the total number of flowering stalks, ignoring seed loss processes.

| Demographic process | Simulated variable | Model | Parameter estimates |
| --- | --- | --- | --- |
| Probability of survival | $Surv \sim Bern(s)$ | GLM | $\text{logit}(s) = -0.024 + 0.456z$ |
| | | GP | $\text{logit}(s) = f_{GP}; r^2 = 0.399; l = 0.978$ |
| Probability of reproduction | $Repr \sim Bern(p_b)$ | GLM | $\text{logit}(p_b) = -5.340 + 2.501z$ |
| | | GP | $\text{logit}(p_b) = f_{GP}; r^2 = 16.87; l = 4.10$ |
| Number of flowering stalks | $Flow \sim Poi(n_f)$ | GLM | $\ln(n_f) = -2.232 + 1.115z$ |
| | | GP | $\ln(n_f) = f_{GP}; r^2 = 7.037; l = 2.628$ |
| Growth for reproducers | $Z' \sim Norm(\mu_r, \sigma_r^2)$ | LM | $\mu_r = 0.656 + 0.767z; \sigma_r^2 = 0.3853$ |
| | | GPR | $\mu_r = f_{GP}; r^2 = 17.68;$<br>$l = 7.093; \sigma_r^2 = 0.3872$ |
| Growth for non-reproducers | $Z' \sim Norm(\mu_n, \sigma_n^2)$ | LM | $\mu_n = 1.284 + 0.528z; \sigma_n^2 = 0.3450$ |
| | | GPR | $\mu_n = f_{GP}; r^2 = 8.117;$<br>$l = 6.354; \sigma_n^2 = 0.3450$ |

Using *D*_gp_ and *D*_glm_, we constructed three types of IPMs (**GLM**, **GP**, and **ABC GP**) to explore the implications of modeling non-linear relationships as linear, and vice versa. When fitting the ABC GP model, we did not borrow the prior knowledge from *M*_gp_ and *M*_glm_ to compute sufficient summary statistics. Instead, we followed the proposed workflow (Workflow 1) to reflect a more generalised modelling process in the real practice. Detailed implementation can be found in Appendix C.

#### 4.1.2 Evaluation Metrics in the Simulation Study

Our analysis focused on evaluating the performance of the fitted IPMs in characterising the entire population dynamics. We calculated key population metrics based on *the stable population growth theory* (Rees & Rose (2002); Ellner & Rees (2006)), such as population growth rate (*λ*), stable state distribution (***ω̅***), stable size distribution for reproductives (***ω̅_rep_***) and relative reproductive value (***γ***). These metrics are widely employed measurements and critical indicators of population dynamics in biological research (e.g. Caswell (2000); Morris & Doak (2002); Rees et al. (2014)). To evaluate the performance of models, we compared the population metrics from the discretized ‘true’ IPMs (generated from *M*_gp_ and *M*_glm_) with those from the estimated IPMs derived from the three modeling strategies.

The deviation of *λ* estimates from the truth was evaluated by Mean Squared Error (MSE), defined as MSE = bias^2^ + variance. The probabilistic metrics ***ω̅*** and ***ω̅_rep_*** were assessed using the Kullback-Leibler (K-L) distance. For ***γ***, which reflects relative contributions, we employed Cosine similarity to measure the angle between vectors, rather than comparing absolute values. Table 3 summarizes the selected population dynamics measurements and their corresponding evaluation metrics.

**Table 3:** For an IPM kernel discretized and evaluated at mesh points (*m*_1_*, …, m_N__m_*), we denote the resulting discretization matrix as K. The matrix K can characterised by its dominant eigenvalue *λ*, alongside its dominant right eigenvector ***ω*** = (*ω*_1_*, …, ω_N__m_*)*^T^* and dominant left eigenvector ***γ***. ***γ*** represents the asymptotic reproductive value associated with individuals in each state. Following standard practice, we normalize it by setting the first entry to 1 to allow for interpretable relative comparisons. Additionally, let ***p_b_*** represent the vector of probabilities of reproduction evaluated at the mesh points. The symbols and denote the Hadamard and dot product between two vectors, respectively. Cosine similarity ranges from 0 to 1, with higher values indicating greater similarity. For MSE and KL divergence, a lower value reflect a closer alignment.

|  | Formula | Meaning | Distance metric |
| --- | --- | --- | --- |
| $\lambda$ | $\lambda$ | Asymptotic population growth rate | MSE |
| $\bar{\omega}$ | $\omega / \sum_1^{N_m} \omega_i$ | Asymptotic population stable state distribution | K-L distance |
| $\gamma$ | $\gamma$ | Asymptotic reproductive value | Cosine similarities |
| $\bar{\omega}_{rep}$ | $(\omega \circ \mathbf{p}_b) / (\omega \cdot \mathbf{p}_b)$ | Asymptotic stable distribution for reproductives | K-L distance |

### 4.2 The real case study

Building on the insights from the simulation case study, we applied the three modelling strategies to real-world datasets of *C. flava* from 2003 to 2011. The models were trained using data from 2003 to 2008, with the final four years reserved for testing. Implementation details are provided in Appendix D.

We evaluated predictive performance by simulating populations with IBMs based on fitted models from the three strategies, focusing on both long-term and 1-step forecasting of demographic trends. Predictive accuracy was assessed through point estimates and 95% prediction intervals, compared to actual observations. Note that, according to the National Oceanic and Atmospheric Administration (NOAA (2013)), 2012 was the warmest year in the historical record for the continental United States from 1895 to 2012. While Gaussian Process models are efficient interpolation machines, we cannot expect them to perform well in such extreme out-of-sample extrapolation effectively.

## 5 Results

### 5.1 The simulation study

#### 5.1.1 Predictive Performance

Figure 2 and Figure 3 provide model fitting results, with detailed numerical results available in Table 8 and 9 in Appendix C. In evaluating the predictive performance of the three model fitting methods on datasets *D*_glm_ and *D*_gp_, the ABC GP method demonstrated more consistent alignment with the true population dynamics. For both datasets, predictions from ABC GP were more closely aligned with the true growth rate *λ* (Figure 2). This trend was particularly evident for dataset *D*_gp_, where ABC GP provided a more accurate reflection of the population growth trend, aligning well with the true *λ* value observed to be less than 1. In addition to predicting *λ*, the ABC GP method also provided reliable estimates of the stable state distribution (***ω̅***) and stable distribution for reproductives (***ω̅_rep_***), as indicated by the minimal mean K-L distance from the true distributions in both datasets (Figure 3). While ABC GP did not achieve the highest similarity scores for the reproductive value (***γ***), its cosine similarity exceeded 0.99, confirming a reliable representation of population dynamics.

**Figure 2:**
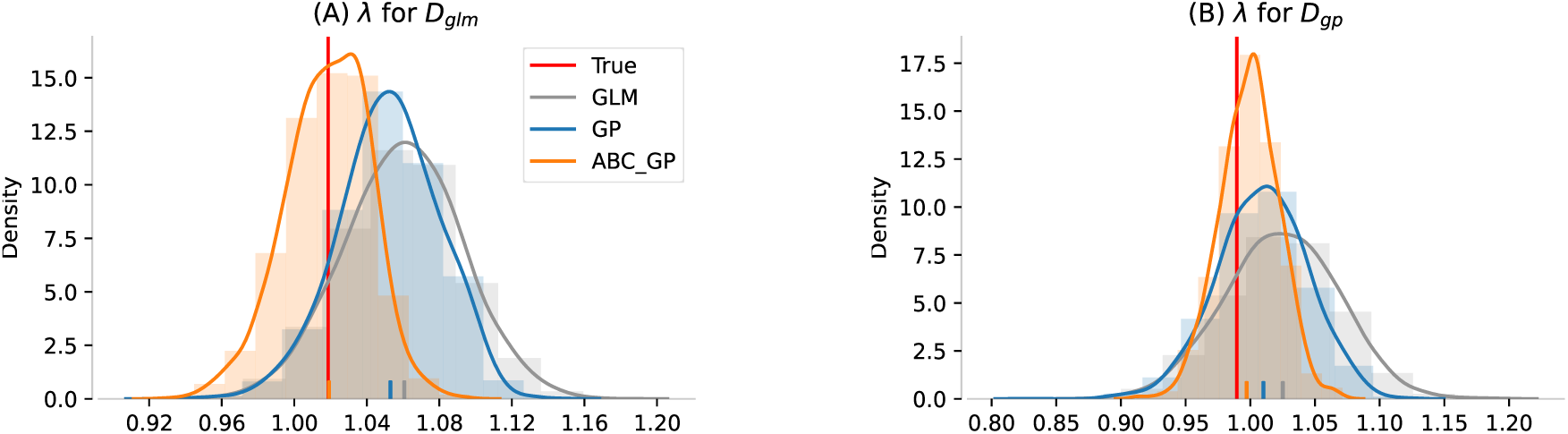
Predictions based on datasets *D*_glm_ and *D*_gp_. Predicted *λ*s, along with their corresponding density histograms (shown as shaded areas), were generated using 5,000 samples draw from the estimated posterior distribution of the three fitting methods. The red line represents the *λ* computed from the true discretized IPM. The pipe (‘|’) in the bottom of each plot marks the corresponding mean.

**Figure 3:**
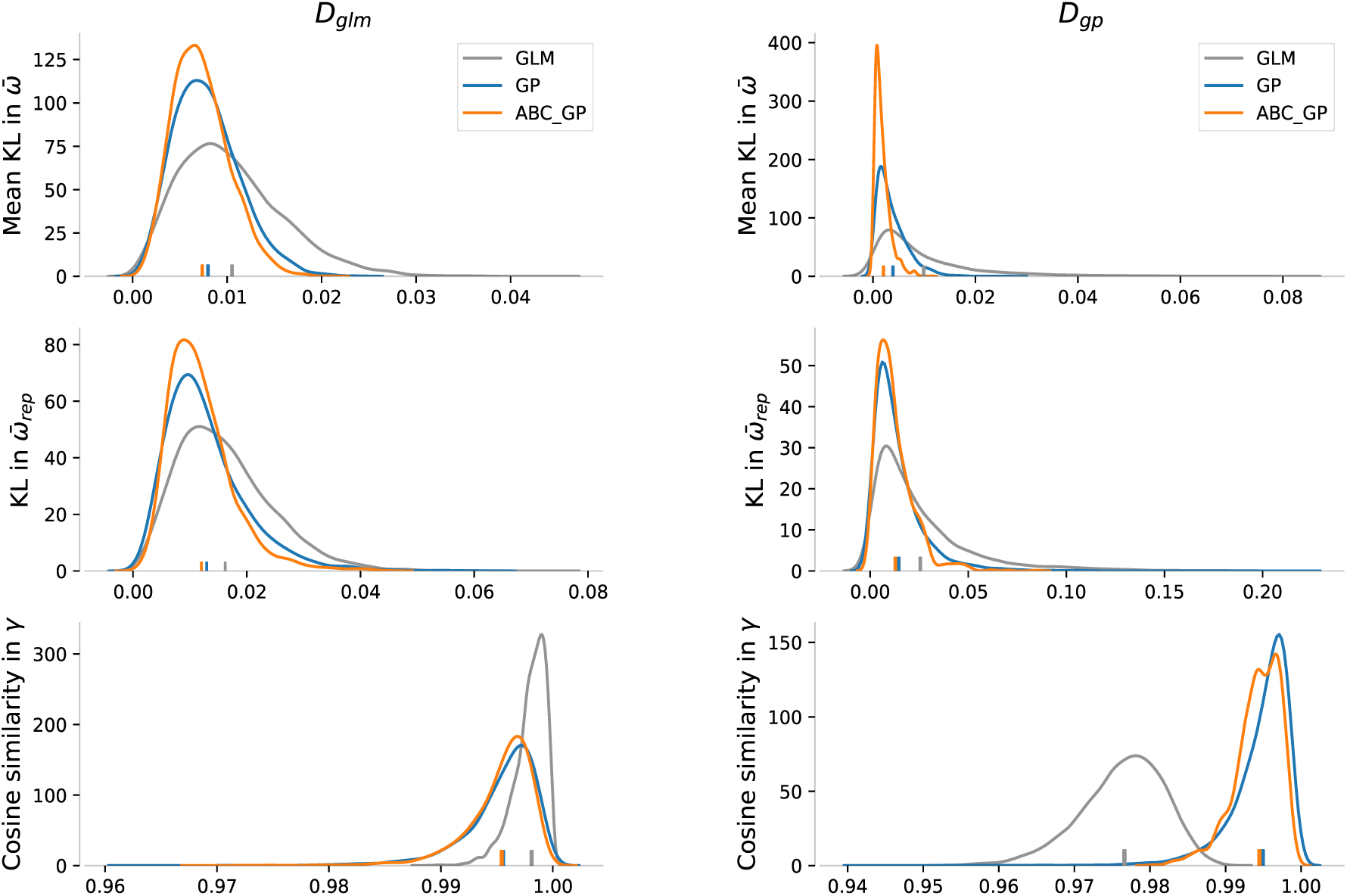
Predictive performance evaluations for the three model fitting methods applied to datasets *D*_glm_ (**left**) and *D*_gp_ (**right**). Different colours represent various settings of the growth model. The pipe (‘|’) in the bottom of each plot marks the corresponding mean.

The GLM method demonstrated reasonable accuracy for dataset *D*_glm_, particularly in estimating *γ*, reflected by the highest cosine similarity scores (the left on third row of Figure 3). However, a noticeable deviation was observed in estimating *λ* (Figure 2), along with inconsistencies in other demographic indicators (Figure 3). These limitations were more noticeable in *D*_gp_, suggesting that GLMs may struggle with the complexity embedded within such datasets.

It is worth noting that while the sample size of *D*_glm_ (562 individuals) is typical for realworld applications, it may not be large enough and remains subject to stochastic effects. To investigate this, a larger dataset *D_glm_*_2_ was generated, comprising information of 4,551 individuals. The newly estimated *λ* from *D_glm_*_2_ demonstrated improved accuracy, closely approximating the true *λ* (Figure 16 in Appendix C), which highlighted the critical role of sample size in GLMs.

Despite the overall strong performance, ABC GP exhibited some deviations in marginal distributions for the vital rate (Figure 4). These discrepancies will be revisited in the Discussion section (Section 6) to clarify their causes and provide guidance for interpretations.

**Figure 4:**
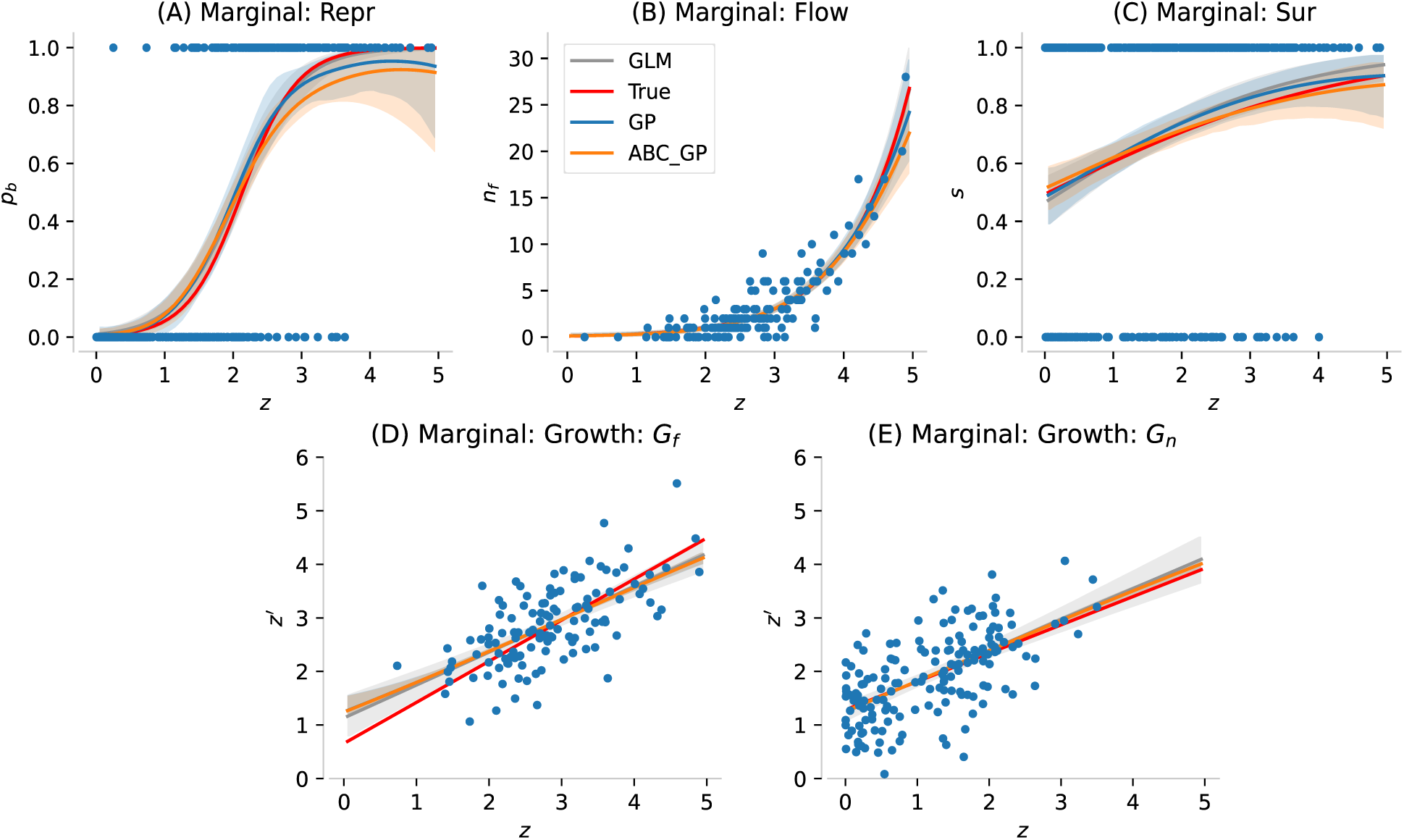
Model fitting results for *D*_glm_. The true and estimated marginal distributions for the five vital rates, complete with their 95% confidence intervals in matching colours and the corresponding scatters data points in blue.

From Figure 2 Left, we saw that the GP and ABC GP produced lower estimates of the population growth rate *λ* compared to the GLM, in *D*_glm_. This difference may be partially explained by the marginal distribution shown in Figure 4 (A), where larger individuals were assigned lower reproduction rates in the GP and ABC GP. To further explore the difference in predicted *λ*, we examined the pairwise joint behaviour among all vital rates (see Figure 18 in Appendix C). In the main text, we focused on survival and reproduction rates, as they showed more visible differences across fitting methods in Figure 4.

Figure 5 (A) compares the survival–reproduction rates across posterior samples from GLM and ABC GP. Since GLM fitted each vital rate independently and deterministically, the implied relationship remained relatively consistent across samples. In contrast, ABC GP captured a range of non-monotonic interactions between survival and reproduction that were not present in GLM, especially in the high-value ranges. As survival increased, reproduction generally rose at first, but in higher survival regions, some samples showed reproduction decreasing, while others continued to increase. Similarly, at high reproduction levels, survival could either decrease or increase across samples.

**Figure 5:**
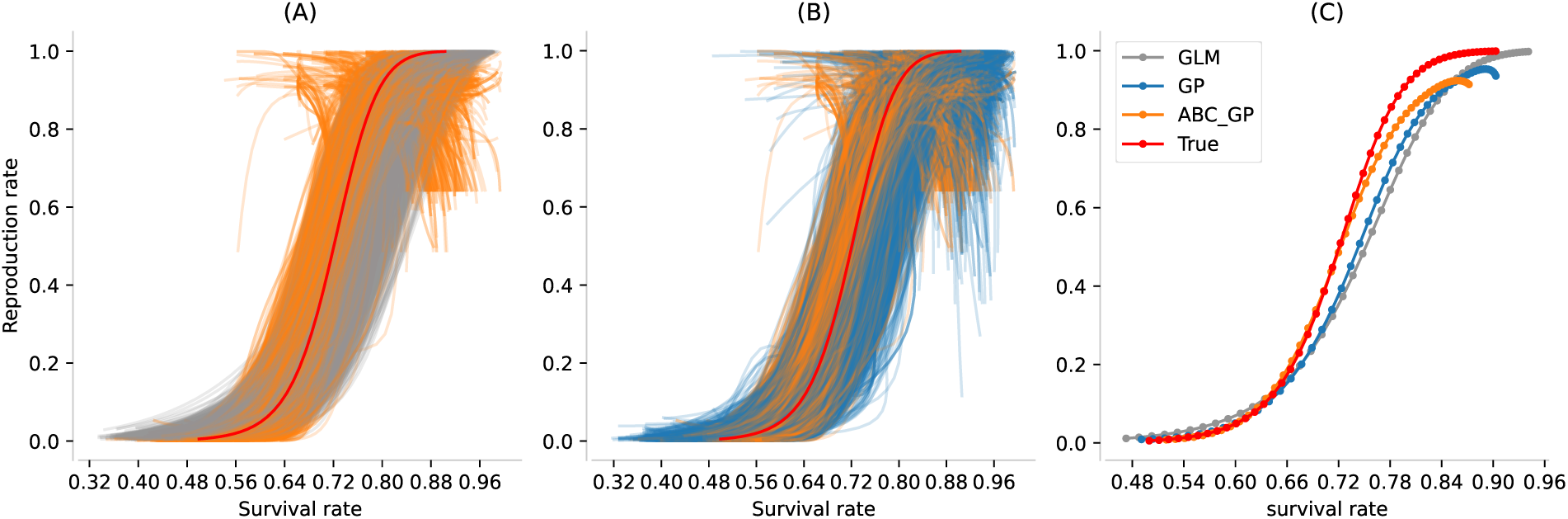
Using the 5,000 posterior samples that generated estimated *λ* in Figure 2 left, we extracted the corresponding survival and reproduction models and evaluated them on 50 mesh points (representing *z* from 0 to 5). The survival–reproduction pairs shown in (A) and (B) were evaluated jointly within each posterior sample at each mesh point, so the joint structure between vital rates was preserved. **(A)** Survival–reproduction pairs from GLM and ABC GP (each line corresponds to a posterior sample evaluated over the mesh points). **(B)** Survival–reproduction pairs from GP and ABC-GP. **(C)** Sample means of survival and reproduction rates at each mesh point, shown for GLM, GP, ABC-GP, and the true model.

Figure 5 (B) shows the same comparison between GP and ABC GP. The ABC step introduced a noticeable shift in the joint patterns. For similar reproduction rates, ABC GP placed more weight on lower survival probabilities. This may help explain why the ABC GP-inferred *λ* is lower in Figure 2 left.

Figure 5 (C) compares the mean survival and reproduction rates at the 50 mesh points across posterior samples. While all models captured the general shapes of the survival and reproduction curves, notable differences appeared in certain regions. For example, both the GLM and GP began to diverge from the truth when survival probabilities exceeded approximately 0.65. In this range, for fixed levels of reproduction, both models tended to overestimate survival rates.

Introducing ABC methods to GP appears to adjust survival-reproduction interaction. However, around a survival probability of 0.72, the ABC GP method also began to deviate from the truth. In this case, again, at fixed levels of reproduction, all models predicted higher survival than the true model.

Interestingly, although the ABC GP model predicts higher survival in these regions, this does not lead to higher overall *λ* values compared to the true model (Figure 2 left). One possible reason may be that samples with both very high survival and very high reproduction are relatively rare in the ABC GP outputs. As shown in the top-right corner of Figure 5(C), the average survival and reproduction rates tended to remain around 0.87 and 0.9, respectively. While the GP results allowed for such joint extremes, these combinations may be filtered out during the ABC step if they fail to reproduce the observed features — for instance, if simultaneous high survival and reproduction are not well supported by the summary statistics.

While such joint extremes are present in the true model, they may be under-represented in the observed data. In the dataset, only 12 out of 562 individuals had size *z* ⩾ 4. This imbalance is also visually reflected in Figure 4 (D), where only a few reproducers fall in *z* ⩾ 4. Together with the positive association between size and survival in Figure 4 (C), this suggests that the limited presence of such large-sized reproducers may make it difficult for the models to learn or retain these joint extremes in survival-reproduction.

### 5.2 The real case study

Figure 6 shows marginal posterior approximations for the hyper-parameters of five vital rate for ABC GP. The hyper-parameters are subject to varying degrees of restrictions based on the chosen summary statistics. For the probability of reproduction (**fec**), all of the six hyperparameters appeared concentrated around a single mode, contrasting with their flat MCMC priors. However, for the Poisson model regarding the number of flowering stalks (**flowp**), the hyper-parameters appeared to be less distinguishable, with no noticeable difference between the MCMC and ABC GP posterior samples in the marginal histograms. Other vital rates also show modifications in MCMC samples due to ABC constraints. The additional information provided by the ABC summary statistics appeared to assign higher weights to certain regions previously having low posterior probability in the MCMC samples, which made the corresponding ABC posterior approximations to be multi-mode, such as the kernel variance for the survival model (**sur kvar**) and the non-breeders’ growth model (**grwnf kvar**), as well as the lengthscale for size and age in the survival model (**sur lsize**, **sur lage**) and the lengthscale for precipitation in the breeders’ growth model (**grwf lprecip**).

**Figure 6:**
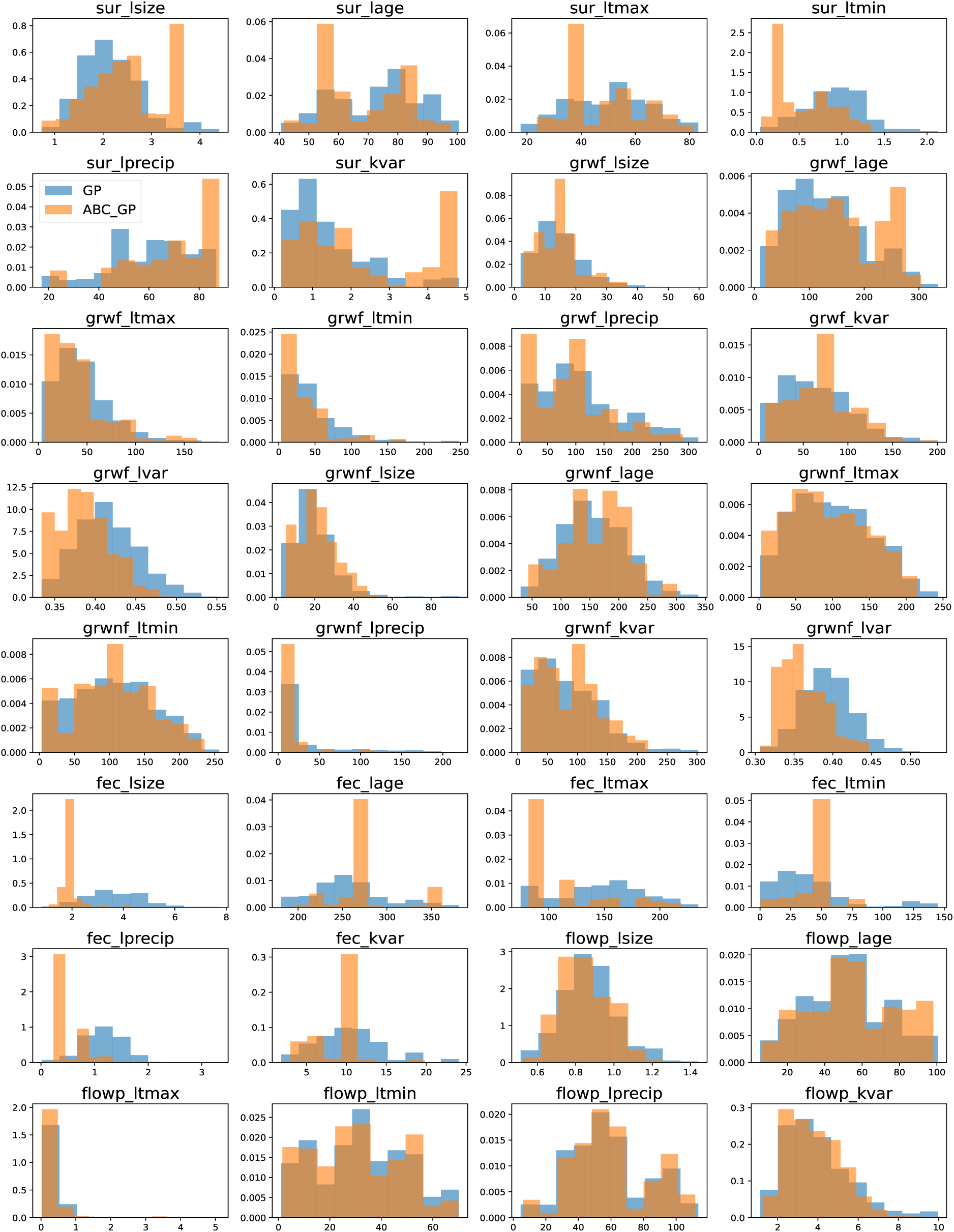
Density histograms of estimated marginal posterior distributions for all hyperparameters under two modelling approaches: GP and ABC_GP.

The impact of ABC restrictions on the marginal posteriors (Figure 6) becomes more evident when integrating individual vital rates into the joint population model (Figure 7 and 8). Median values were used as point estimates to reduce outlier impacts in IBM simulations. A side-by-side comparison with GLM and GP methods revealed that, GLM-based models tended to produce larger absolute errors in predicted population size (see Figure 7). In certain cases, such as the 1-step forecast in Figure 8, GLM predictions showed notable deviations, with trends often contradictory to actual observations. While GP methods also showed some deviations (particularly for 2009), switching to ABC GP led to improvements in accuracy for both long-term and 1-step forecasts.

**Figure 7:**
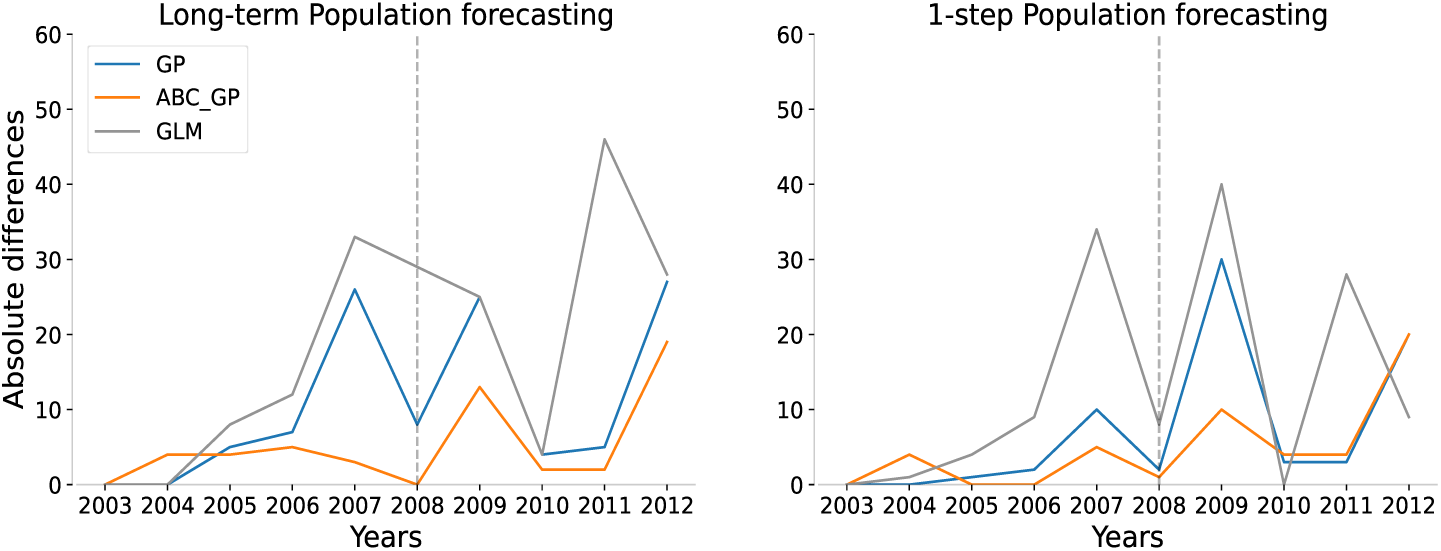
Absolute differences between the observed and predicted population size (medians of the predictions). The models were trained using five datasets from years 2003-2004 to 2007-2008 (before the grey dashed vertical line). For **long-term** forecasting, the model used the initial population structure from Year 2003 as the starting point and generated predictions based on simulated populations from the previous year. For example, the predicted population size for 2005 was based on the simulated population for 2004 (which was generated using the true observed population for 2003). For **1-step** forecasting, the model used the true observed population structure from the previous year to generate predictions. That is, for example, to make a prediction for 2005, the true structure for 2004 was plugged into the IBM.

**Figure 8:**
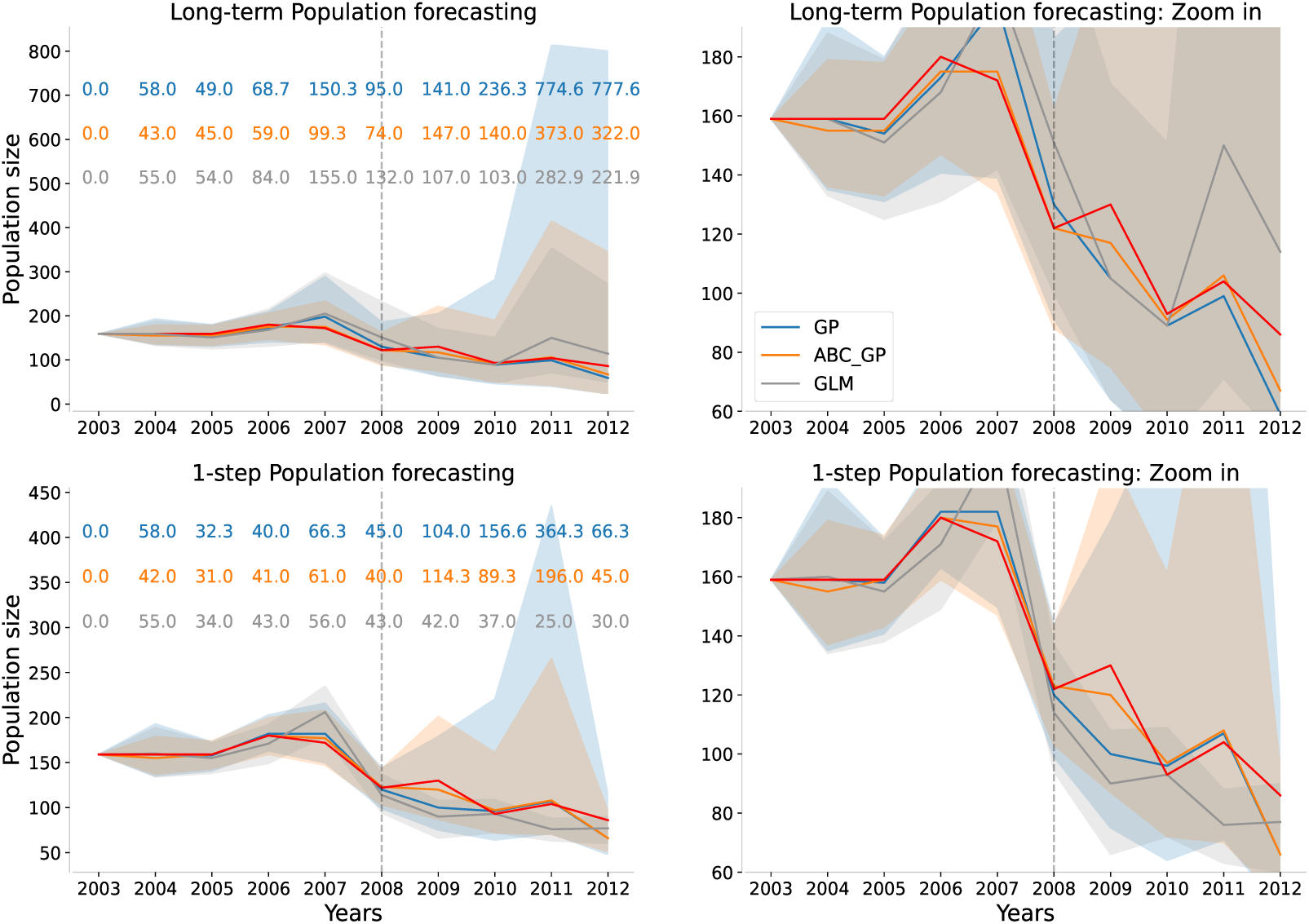
Two types of population size predictions, along with their corresponding 95% PIs (shaded areas), were generated using GP, ABC GP and GLM samples through 5,000 IBM simulations. The blue predictions were based on MCMC samples, while the orange and grey ones were based on ABC GP and GLM posterior samples, respectively. The widths of the corresponding 95% PIs are indicated by their respective colours on the left-hand side of the figure. The point estimations are represented by solid lines in their corresponding colours. The red line represents the true observed population size.

For long-term population forecasting, prediction intervals (PIs) of all three model types remain narrow during training years and widen in subsequent periods, generally covering the observed population sizes. However, GP models exhibit broader PIs compared to the other models.

More details can be illustrated by PIs for 1-step predictions, which reduce the influence of cumulative demographic randomness present in long-term forecasts. GP models show a similar pattern as in long-term predictions, but the width of the PI generated by GLMs remains consistently narrow, regardless of whether applied to training or testing datasets.

Although GLMs exhibited narrow PIs, their reliability appears limited. As detailed in Figure 9, focusing on the annual population growth rate, we observed that the actual growth rates frequently fell outside the GLM’s 95% PIs, indicating a mismatch even for the training year 2006-2007. In contrast, the ABC GP method not only covers the true growth rates but also adjusts its estimates to align with observed changes. Among the three models considered,

**Figure 9:**
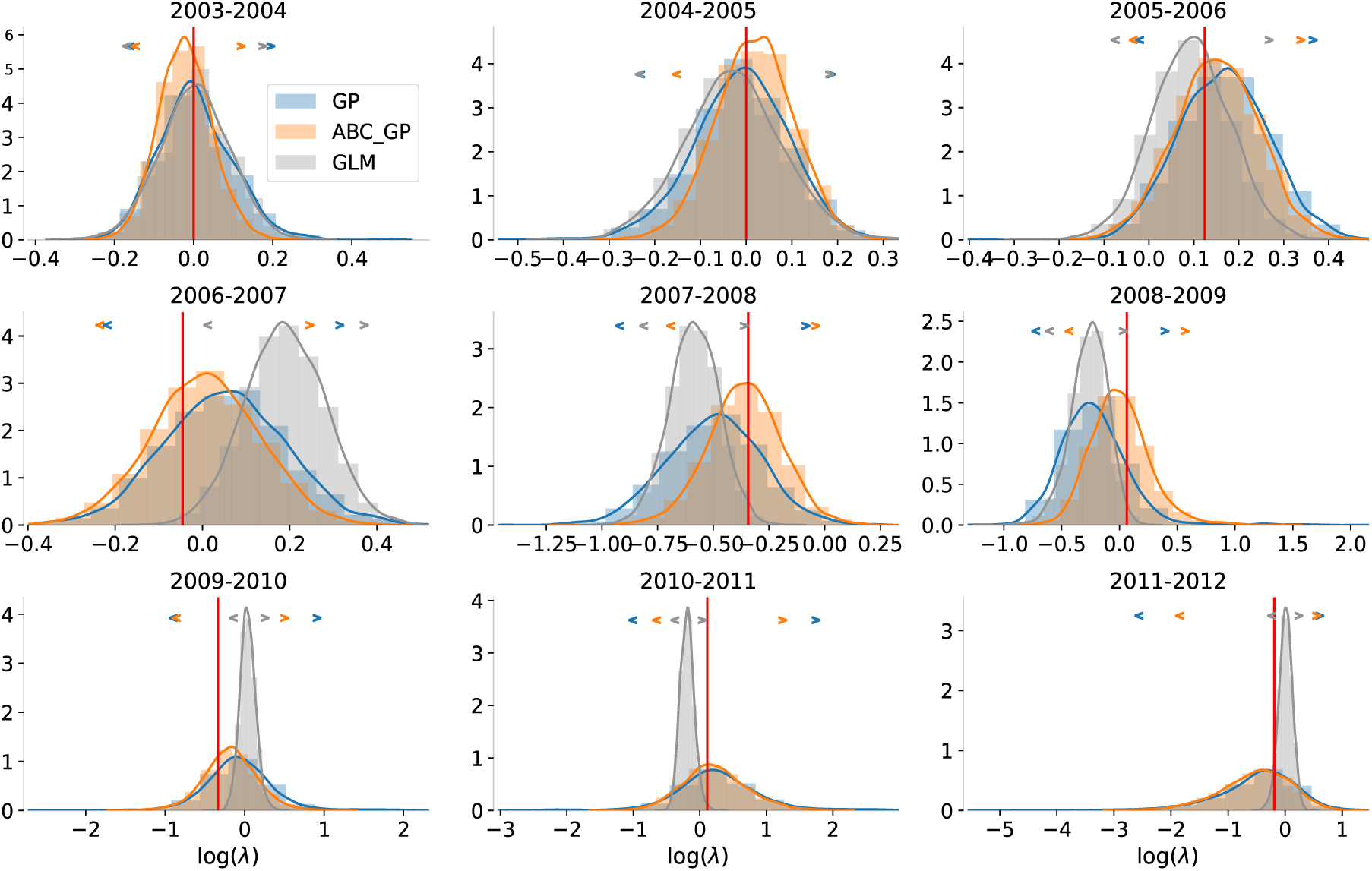
Histograms represent the density of logarithmic simulated population growth rates generated through **1-step** forecasting. The associated 95% prediction intervals are denoted by angle brackets, each in its corresponding colour, positioned at the top of each histogram. The red vertical line represents the true observed population growth rate.

## 6 Discussion

In this work, we introduced a novel methodology, ABC GP IPM, aimed at enhancing the accuracy and flexibility of IPMs. In the simulation study, by integrating population-level information, ABC GP IPM showed notable improvements over traditional GLM-based IPMs across multiple population metrics commonly used in biological research. In the real-world case study, the method maintained accuracy in point estimates while providing a more honest representation of uncertainties in predictions, particularly when the true relationship between predictors and the response was complex or not well defined. In Zhu et al. (2025) we examine the efficiency of fitting this novel IPM through the ABC-PMC algorithm at greater length, revealing these improvements were achieved progressively across sampling iterations.

A key factor contributing to these improvements is the interaction between GP models and ABC. As Bayesian non-parametric models, GP models can theoretically accommodate an infinite number of ‘parameters’. However, with real-world finite datasets, they only utilise a finite subset of these ‘parameters’, with the remainder being marginalised out (Orbanz & Teh (2010)). Within the ABC framework, the summary statistics serve as additional data points, thereby expanding the existing dataset available to the GP models. As more information is incorporated, the ABC GP IPM adaptively adjust their complexity to better reflect the increased volume of information.

This adaptive adjustment enables ABC GP IPM to increase model complexity without fundamentally altering the underlying framework, maintaining its data-driven nature while requiring no additional assumptions beyond the observed data. Despite its flexibility, this approach maintains clarity and interpretability. In addition, ABC GP IPM delivers these improved outcomes using the same datasets as standard IPMs, so no additional data are needed. This makes it convenient for studies that have previously used traditional methods. Its adaptability suggests potential applicability across various demographic studies. We conclude with some final remarks regarding the proper interpretation of the approach and potential concerns.

### Why is the overall performance Good, but the marginal distribution Poor?

The adoption of ABC method shifts the focus from solely maximising likelihood to, additionally, minimising the discrepancy between model predictions and observed population data, through carefully selected summary statistics. Under this dual focus, while the population-level projections often align well with observed data compared to traditional IPMs (Figure 2)), the accuracy of individual vital rate distributions may be compromised (Figure 4).

A natural question then arises: if our primary interest lies in individual vital rates, would not such biased marginal results be misleading? The answer fundamentally depends on the research objectives. ABC introduces additional complexity to IPMs by incorporating population-level feedback. As a result, models calibrated to capture joint population dynamics, may not provide reliable estimates for individual vital rates. This is because, not all population-level insights can directly inform specific vital rates.

In our simulations, where life cycle components are clearly defined, we are able to accurately project the high-dimensional population model onto the ‘correct’ dimensions for individual vital rates. However, this level of clarity is often lacking in real-world settings, where life cycle structures might be partially obscured or unknown. In such cases, projecting a complex population model onto misaligned dimensions can lead to less informative or even misleading interpretations. This situation is akin to viewing a cylindrical object from different angles, where each angle offers a distinct 2D projection, leading to varying interpretations of the original 3D object.

Therefore, again, it is crucial to choose the modeling approach based on specific research objectives. If the focus is on understanding population-level dynamics, the ABC approach provides a more cohesive representation. However, if estimation of individual vital rates is the primary goal, traditional likelihood-based methods may be more appropriate.

### ABC’s role in revealing interactions detected among vital rates

It is important to clarify that our method’s capacity to detect interactions among vital rates should be viewed as a bonus from employing ABC in IPMs, rather than a dedicated tool for studying the joint behaviours. Indeed, correlations in vital rates are inherent in IPM; for example, all vital rate models are often depend on the shared state variables (Fung et al. (2022)). Here, ABC GP IPM amplifies these interactions by incorporating population-level feedback during model assembling.

The method leverages ABC methods and population-level information to narrow down under-determined vital models, retaining only the joint distributions that satisfy specific criteria. As these inherent interactions are not systematically formulated, they might be tricky to incorporate in studies where the sensitivity of these interactions can substantially impact uncertainty of model predictions (Fieberg & Ellner (2001)), since varying these interactions can be difficult (Ellner et al. (2016)).

While our approach can not directly explain how interactions shape life-history evolution, it equips IPMs to better ‘mirror’ observations via ABC. This could open doors to studying interactions from a fresh perspective. By successfully replicating observed behaviours and generating predictions, these models can guide researchers in designing new experiments or identifying the type of data required to further test these mechanisms (Farrell et al. (2021)).

### Managing background knowledge integrated in ABC GP IPM

In the Introduction, we proposed that our model offers scientists greater flexibility in integrating their expertise with IPMs. This claim, however, may spark discussion, as prior ABC research suggests that unsuitable summary statistics can bias inferences (Prangle (2017)). This point is also exemplified in Zhu et al. (2025) where an unreliable inference result was generated due to our over-reliance on an unsuitable summary statistic in a Poisson model. This conclusion was drawn on the premise that the true underlying model was Poisson.

However, in real-world practice, it is critical to ask: “who genuinely knows the true model that generated the observed data?” If our background knowledge suggests greater confidence in the chosen summary statistic over the Poisson assumption, it implies that the Poisson model, not the summary statistic, may be the one ‘mis-matched’. In this case, the derived ABC posterior approximation can be therefore treated as a set of parametric values most likely to align with our intended results under this (potentially) unsuitable Poisson model. In reality, where the true model is unknown, an observed statistic should indeed hold potential to serve as a valid summary statistic in ABC, especially if expert knowledge supports its relevance.

The debate, concerning the appropriateness of the model or summary statistics, may provide another reason for applying ABC to non-parametric models, like in our case with GP models. Unlike parametric models, GPs do not assume a fixed form for underlying probability functions.

While our methods offer a flexible way to integrate users’ own background knowledge into IPMs, it is crucial to control the volume of content added to avoid falling into the curse of dimensionality. We suggest constructing a candidate list comprising all aspects of interest and then employing a systematic approach to pick out the final statistics, as demonstrated in our study. Some statistics in the candidate list may bring overlapping information, causing certain statistics to become less informative when others are present. Techniques like Principal Component Analysis can be employed to help, but the resultant statistics might not be as interpretable as their original counterparts.

### Lacking sufficiency in non-parametric modelling

On a different note, when applying ABC and Bayesian non-parametric models into combined models, the concept of sufficient statistics, in the usual sense of the term, is less clear and might not be applicable given our current knowledge. Non-parametric models do not make strong assumptions about the data distribution or the functional form of the model, making it challenging to identify a set of statistics that can summarise all the relevant information in the data. These models are designed to capture as much information as possible from the data, often utilising the entire dataset for predictions. In this case, the ‘statistics’ calculated from the data should essentially be the data points themselves. As such, while there might be ways to summarize the data in non-parametric problems, but should not referred to as “sufficient statistics” in the same sense as in parametric models. Additional theoretical work is required to clarify how ABC can be applied to non-parametric models, particularly considering the lack of a clear concept of sufficiency in these models, as compared to their parametric counterparts.

## Acknowledgements

For the purpose of Open Access, the author has applied a CC BY public copyright licence to any Author Accepted Manuscript version arising from this submission.

## A The ABC-PMC algorithm

**Table 4:** The description of notations used in Algorithm 1.

| Notation | Value | Description |
| --- | --- | --- |
| $\mathcal{M}$ | | IPM |
| $\top$ | | Transpose |
| $C$ | 10 | The total number of simulation steps for ABC-PMC (apart from the initialisation) |
| $c$ | | Current simulation step; $c = 0$ for initialisation |
| $N_c$ | | The number of accepted particles required for the simulation step $c$ |
| $\alpha_c$ | | The quantile employed to decide the acceptance region for the next simulation $c + 1$. |
| $V$ | 5 | The number of different vital rates in the IPM |
| $v$ | | A realisation of a discrete uniformly distributed random variable indicating the current vital rate. |
| $M$ | 5000 | The MCMC sample size for each of the five vital rates. So, there are $M^5$ different combinations in total. |
| $m_v$ | | Given the current vital rate $v$, $m_v$ is a realisation of a discrete uniformly distributed random variable representing $m_v$th MCMC sample for $v$th vital rate. |
| $\theta$ | | A particle: a vector including parameters for all of the five vital rates. |
| $\theta_c^i$ | | $i$th particle at $c$th simulation step |
| $\tilde{\theta}$ | | An intermediate particle |
| $\tilde{\theta}_v$ | | A vector only includes $v$th vital rate's parameters. So, $\tilde{\theta}^\top = (\tilde{\theta}_1^\top, \dots, \tilde{\theta}_5^\top)$. |
| $\tilde{\theta}^*$ | | A perturbed intermediate particle |
| $\pi(\theta)$ | | Non-informative prior for $\theta$, which is the joint distribution for $m_1, \dots, m_5$. |
| $\omega_c^i$ | | Unnormalised weight for $\theta_c^i$ |
| $w_c^i$ | | Normalised weight for $\theta_c^i$ |
| $y^*$ | | A simulated dataset |
| $y_{obs}$ | | The observed dataset |
| $\mathcal{d}_s$ | | Dimension of $\mathbf{S}_c^i$ and $\tilde{\mathbf{S}}^*$ |
| $\mathbf{S}_c^i$ | | $\mathbf{S}_c^i = ({}^1S_c^i, {}^2S_c^i, {}^3S_c^i, {}^4S_c^i, \dots, {}^{d_s}S_c^i)$, where $({}^jS_c^i)$ is $j$th summary statistics for $i$th particle at step $c$. |
| $\tilde{\mathbf{S}}^*$ | | A vector of summary statistics produced by $\tilde{\theta}^*$, $\tilde{\mathbf{S}}^* = ({}^1\tilde{S}^*, \dots, {}^{d_s}\tilde{S}^*)$, (i.e. the distances between a simulated and the observed dataset through the list of selected combinations of summary statistics and distance metrics $\tilde{\mathbf{S}}^* \in \mathbb{R}^{d_s}$). |
| $\sigma_c$ | | A vector of median absolute deviations (MADs) for the $\mathcal{d}_s$ dimensional summary statistics, $\sigma_c = ({}^1\sigma_c, \dots, {}^{d_s}\sigma_c)$, where ${}^i\sigma_{c-1}$ is the estimated MAD (based on $c - 1$th generation of particles) for the $i$th summary statistic ${}^i\tilde{S}^*$. |
| $d_c^i$ | | $d_c^i = (\sum_{j=1}^{d_s} ({}^jS_c^i / {}^j\sigma_c)^2)^{1/2}$, which represent the ‘universal’ distance between $y_{obs}$ and a simulated dataset produced by $\theta_c^i$. |
| $\tilde{d}^*$ | | $\tilde{d}^* = (\sum_{j=1}^{d_s} ({}^j\tilde{S}^* / {}^j\sigma_{c-1})^2)^{1/2}$, the ‘universal’ distance between the observed and a simulated dataset. |
| $\delta_c$ | | $\alpha_c$th quantile of $\{d_c^i\}_{1 \leq i \leq N_c}$; the quantile-based acceptance threshold at step $c$. |
| $\mathbb{1}_\tau$ | | An indicator function, $\mathbb{1}_\tau = 1$ if the statement $\tau$ is true; $\mathbb{1}_\tau = 0$ otherwise. |
| $a \rightarrow b$ | | This is a statement about whether it is possible to transit from state $a$ to state $b$ by the perturbation kernel at line 15 in Algorithm 1. |

Algorithm 1: ABC-PMC (see Table 4 for the descriptions of the notations)

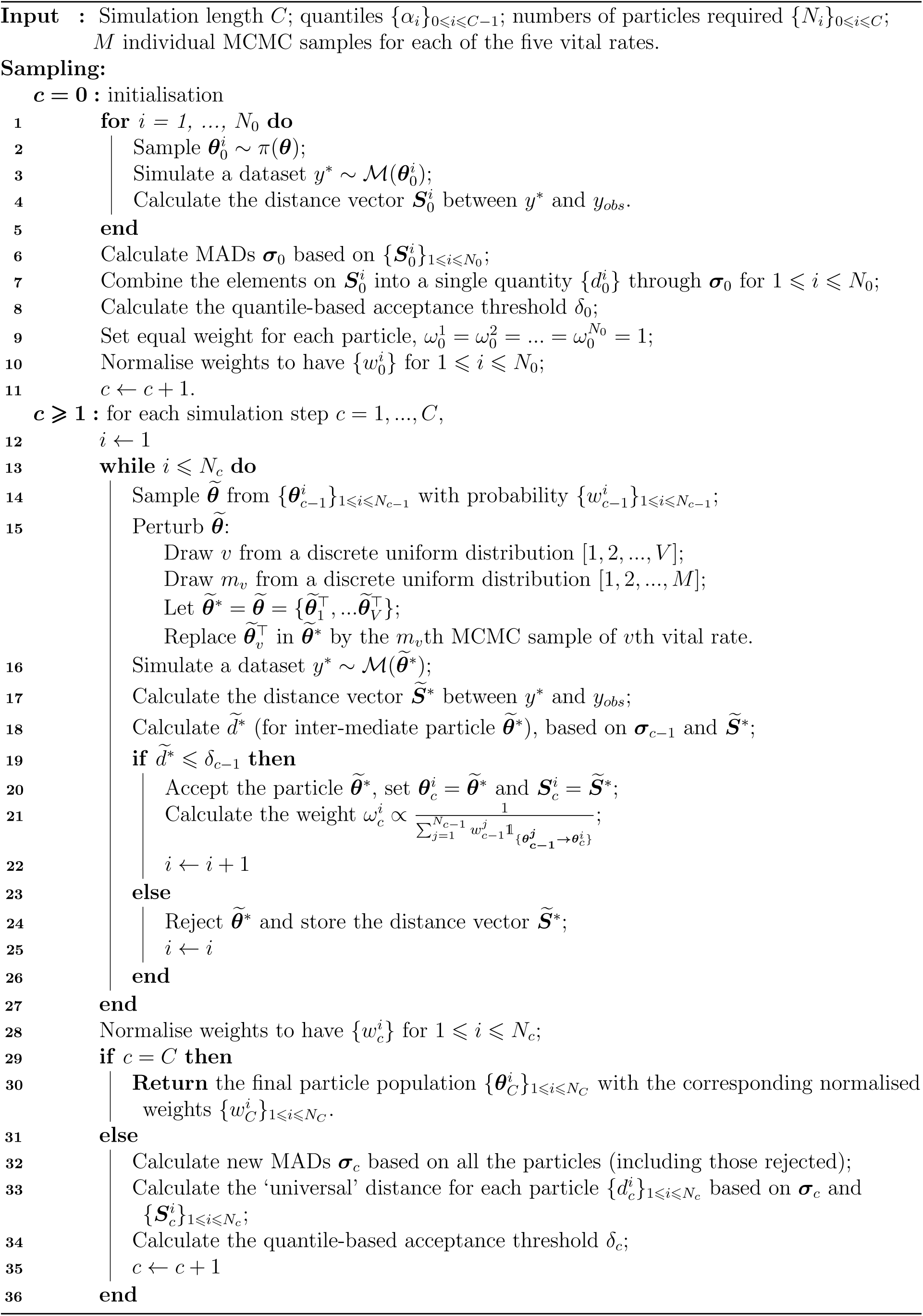

## B Summary statistics and distance metrics

### B.1 A full description for candidate summary statistics and distance metrics

To calculate most of the candidate summary statistics, individuals are first grouped based on their size, either their current size or their size in the following year (denoted as sizeNext). Twelve size/sizeNext groups are defined according to the logarithm of size/sizeNext, using the following bin edges: [0, 0.3, 0.6, 0.9, 1.2, 1.5, 1.8, 2.1, 2.5, 3, 3.5, 4*, >* 4]. For many candidate summary statistics, their meanings are self-explanatory by their names. However, we highlight the following summary statistics that may require further clarification:

- **The empirical population distribution in sizeNext groups:**the counts of individuals in the corresponding groups divided by the total number of individual in the entire population.
- **Average number of flowering stalks/breeders/survivors in size groups:** the counts of flowering stalks/breeders/survivors in the corresponding groups divided by the total number of individuals in the corresponding group.
- **breeders’/non-breeders’ sizeNext distribution:** the counts of breeders/non-breeders in the corresponding groups divided by the total number of individual in the entire population.
- **breeders’ and non-breeders’ survival distribution:** individuals are divided into four groups based on whether they were breeders in the past year and whether they survived in the past year.

Some summary statistics are constructed from frequencies, while others are based on distributions. Accordingly, we employed different distance metrics based on their attributes. The complete list of employed distance metrics can be found below. Let *P* and *Q* denote two (empirical) discrete distributions, where *p_i_* and *q_i_* are the *ith* element of distributions *P* and *Q*, respectively. Similarly, let *U* and *V* represent two sets of values, with *u_i_* and *v_i_* denoting their *i*th elements:

1. **Bhattacharyya distance:** is a measure of similarity between two probability distributions. For discrete probability distributions, the distance is defined as 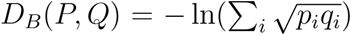.
2. **Hellinger distance:** is another measure of similarity between two probability distributions. It is related to the Bhattacharyya distance and is defined as 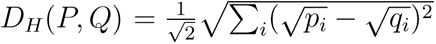.
3. **Earth Mover’s Distance (EMD)** (also known as the Kantorovich-Rubinstein norm or Wasserstein distance): is a measure of the distance between two probability distributions, interpreted as the minimum amount of “work” necessary needed to transform one distribution into the other. The EMD is defined as: 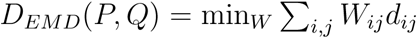 where, *W* is a flow matrix representing the optimal transportation plan, and *d_i_j* is the difference between the elements *i* and *j* of the distributions. Here, “work” is calculated by multiplying the amount of weight that must be relocated by the distance it must travel.
4. **Kullback-Leibler (K-L) distance** (also known as relative entropy): is a measure of how one probability distribution is different from a second. It is defined as 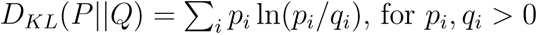.
5. **Hilbert projective metric:** a distance function defined on the positive cone of a real Hilbert space. It defined as *D_H_* (*P, Q*) = log(max*_i_*(*p_i_/q_i_*)) + log(max*_i_*(*q_i_/p_i_*)), for *p_i_, q_i_ >* 0.
6. **Chi-square (***χ*^2^**) distance metric:**The chi-square distance is a measure of dissimilarity between two sets of values and is often used in comparing histograms. It is defined as *χ*^2^(*U, V*) = Σ*_i_*(*u_i_* − *v_i_*)^2^*/*(*v_i_*).
7. **Sum of squared differences (SSD):** a common metric used to measure the dissimilarity between two sets of values. It is defined as: *SSD*(*U, V*) = Σ*_i_*(*u_i_* − *v_i_*)^2^.
8. **Cosine similarities:** a common metric used to measure the dissimilarity between two vectors. It is defined as: cosine similarity 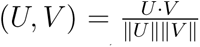. Here, *U · V* is the dot product of vectors. *∥U∥* and *∥V∥* are the magnitudes (or lengths) of the vectors. The cosine similarity ranges from −1 to 1, where 1 indicates that the vectors are identical, −1 indicates that they are completely opposite, and 0 indicates orthogonality (no similarity).

All calculations were implemented using self-developed Python code, except for the EMD, which was computed using the wasserstein_distance function from package scipy. Both the K-L distance and the Hilbert projective metric require *p_i_* and *q_i_* to be greater than 0. Our corresponding functions were therefore modified to detect if *p_i_ <* 0 or *q_i_ <* 0 occurred. If either condition occurred, the functions would automatically merge the neighbouring cells in both *P* and *Q* to ensure comparability, until *p_i_ >* 0 and *q_i_ >* 0 for all *i*. A similar strategy was applied to the *χ*^2^ distance metric to ensure that the frequencies in the expected values were greater than 5.

### B.2 A note on using NLPP to define similarity between two GP models

GP models are highly flexible. When perturbing a single hyperparameter, the corresponding effect is highly dependent on the values of the other hyperparameters.

To explore this, we first use a toy dataset to perform Bayesian inference on a GP regression model predicting breeders’ growth (*G_f_*). Two representative samples are selected from the posterior, labelled as **M1** and **M2**. Both samples correspond to the same GP regression model and exhibit similar Negative Log-Likelihood (NLL) scores, indicating comparable performance in data fitting (we employed non-informative priors in our study, thus the NLPP closely approximates the NLL (NLPP NLL), due to the minimal impact of the prior). Next, we implement standard perturbation to the hyperparameters in **M1** and **M2**, by adjusting one hyperparameter at a time while keeping the others fixed. The predicted mean and variance for the models, before and after the perturbation, are shown in Figures 10 and 11 respectively.

**Table 5:** The values of the hyper-parameters of two GP regression models in Figure 10 and Figure 11 (growth model for breeding individuals). Models producing Plot (b) and (d) were produced by perturbing the lengthscale for the log size with the same absolute amount of noise, −1, on (a) and (c) respectively.

| Plot | Kernel | NLL | $n_s$ | Scale $r^2$ | ARD | |
| --- | --- | --- | --- | --- | --- | --- |
|  |  |  |  |  | Size | Age |
| (a) | Gau | 54.02 | 0.3243 | 8.533 | $l_1 = 7.098$ | $l_2 = 18.31$ |
| (b) | Gau | 53.94 | 0.3243 | 8.533 | $l_1 = 6.098$ | $l_2 = 18.31$ |
| (c) | Gau | 54.52 | 0.2952 | 4.480 | $l_1 = 2.233$ | $l_2 = 5.531$ |
| (d) | Gau | 56.51 | 0.2952 | 4.480 | $l_1 = 1.233$ | $l_2 = 5.531$ |

**Figure 10:**
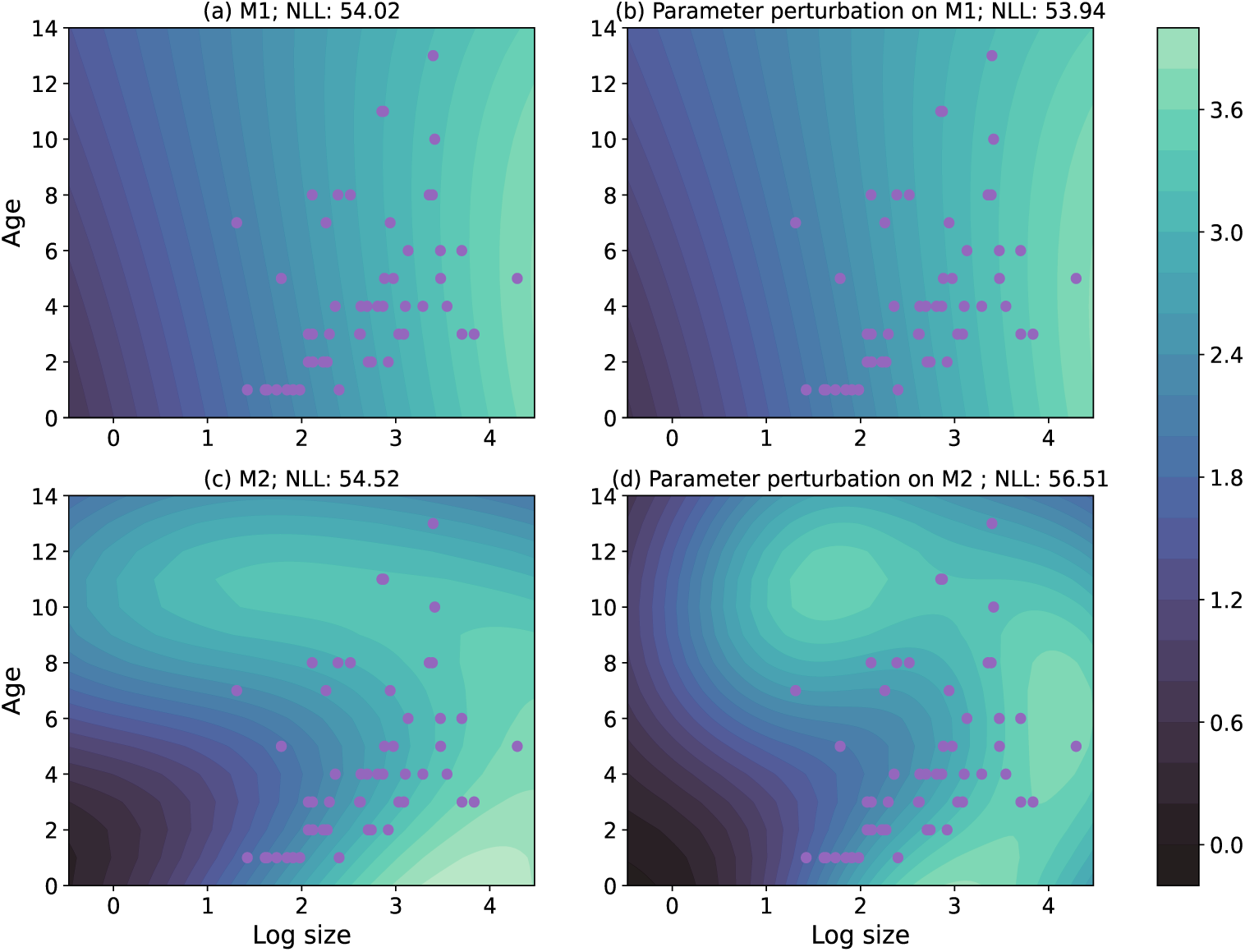
Given the current age (*y*-axis) logged size (*x*-axis), the contour plots represent the predicted mean for logged size in the next year. Predictions were made through Gaussian process models with hyper-parameters shown in Table 5. Plot (b) and (d) were produced by perturbing the lengthscale for the log size with the same **absolute** amount of noise on (a) and (c) respectively. Purple dots represent for the values of covariates for the training data points.

**Figure 11:**
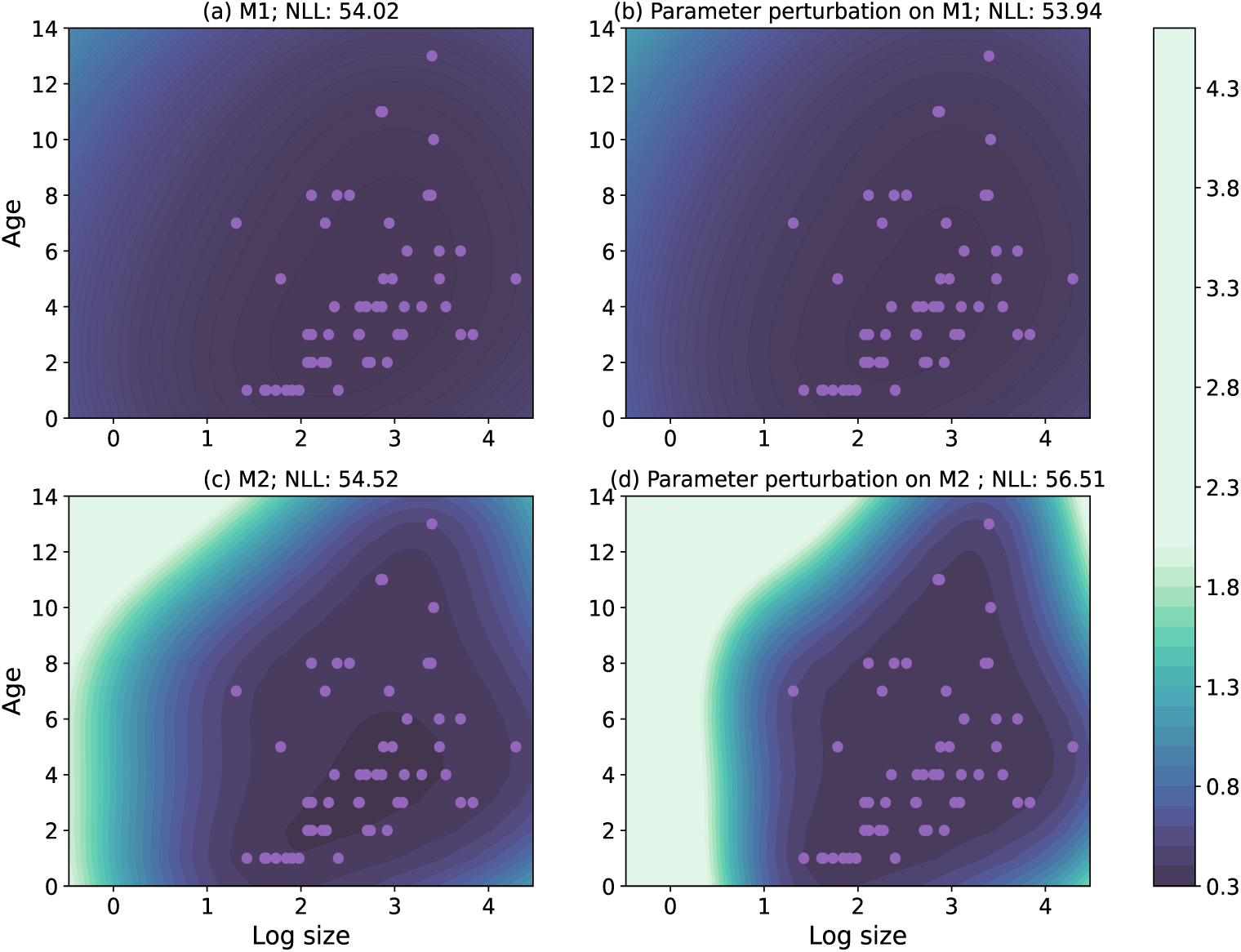
Contour plots for the predicted variance corresponding to the predicted mean shown in Figure 10.

When the same hyperparameter of **M1** and **M2** are perturbed with the same absolute amount, the contour plots of the predicted mean/variance for **M1** show no obvious difference before and after the perturbation (see plot (a) and (b) in Figure 10/11), whereas **M2** produces dramatic changes. Its predicted mean begin to show multiple peaks (plot (c) and (d) in Figure 10), and the predicted variance fall much more sharply as the logged size increased (plot (c) and (d) in Figure 11). When quantifying the effect of the perturbations, we observe that the NLL score for **M1** decreases from 54.02 by 0.08 to 53.94. However, for **M2**, the score changes from 54.52 to 56.51, with a difference of +1.99. Similar results can be also found in Figure 14 and Figure 15 in the end of this notes, which considers the response to the effects of proportional perturbations instead.

The impact of perturbations varies due to interactions among hyperparameters in GP kernels. These interactions become even more complex as the input dimension increases.

Let us revise the interpretation first. The Gaussian kernel (Gau) with hyper-parameter output variance *r*^2^ and lengthscales *l* measures the similarity about inputs *x_i_* and *x_j_* through

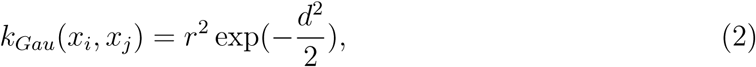

Where 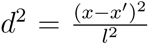 for 1-dimensional inputs. Given a kernel *k_Gau_*, the entries of covariance matrix *K* used in a GP model is calculated through *K_i,j_*= *k_Gau_*(*x_i_, x_j_*) for *i* and *j*th inputs. The *k_Gau_*takes two parameters.

**Figure 12:**
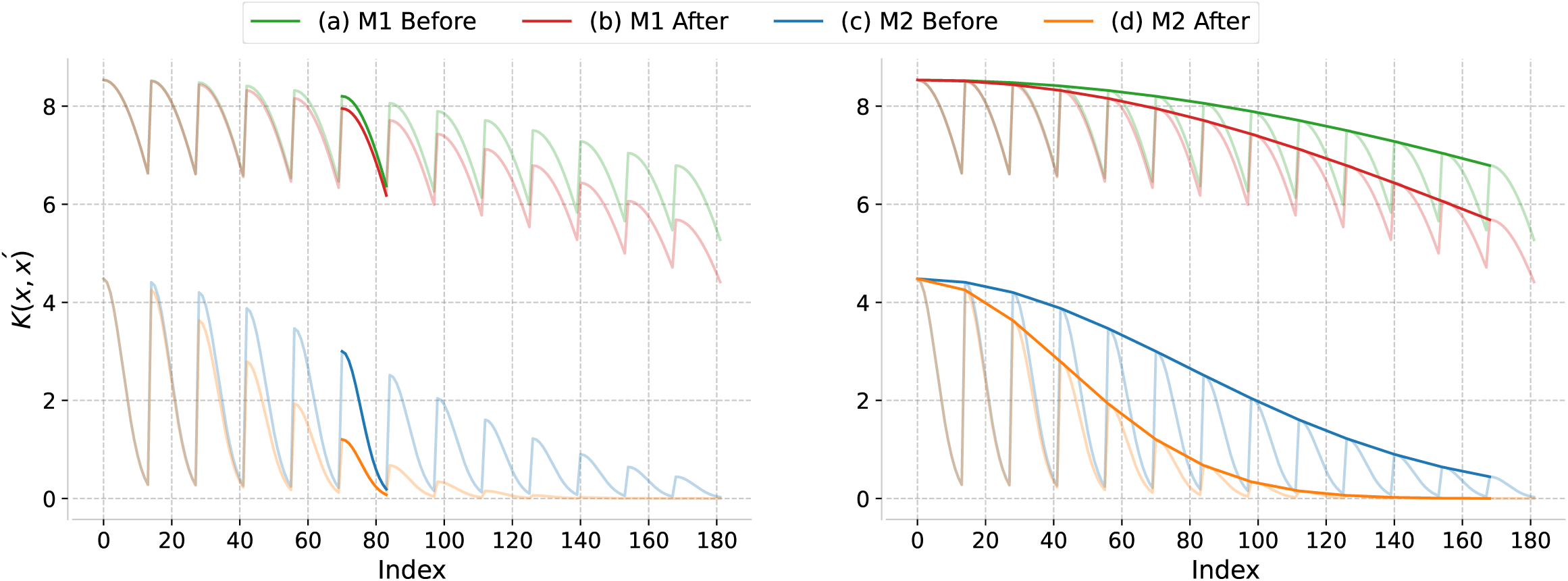
The plots display samples obtained from GP models with kernel function *k_Gau_*. The left plot displays the samples obtained by varying the scale factor *r* from 1 to 6 while fixing the lengthscale *l* at 1. Conversely, the right panel illustrates the effect of increasing *l* from 1 to 6, while keeping *r* fixed at 1.

- The scale factor *r*^2^ controls the average distance of the results away from their mean. For example, looking at Figure 12 left, the ‘amplitude’ increases when raising *r* from 1 to 6. Almost every GP kernel has this scale hyper-parameter *r*^2^ to speak for the output variance.
- The lengthscale *l*^2^ is used to adjust the strength of correlation between data points (as measured by the squared Euclidean distance). From Figure 12 right, a shorter lengthscale would bring us a “more wiggly” curve when keeping *r*^2^ fixed. As a smaller length scale implies the data points far away impact less to the current point, then results a more rapidly varying function. A extreme large value in *l* therefore corresponds to a linear covariate.

Back to the exploration of **M1** and **M2**, we have the case of 2-dimensional inputs, e.g. ***x_i_*** = (*x*_1_*_,i_, x*_2_*_,i_*)^⊤^. With automatic relevance determination (ARD) in GP models, we can introduce a separate lengthscale for each dimension. The kernel function *k_Gau_* with this ARD setting can be expressed in two forms:

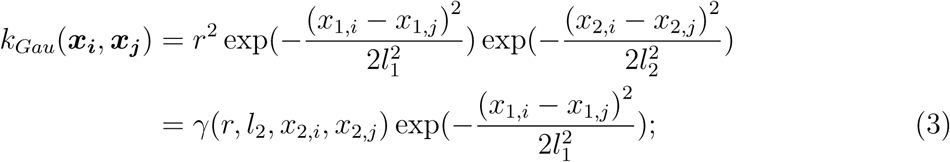

or,

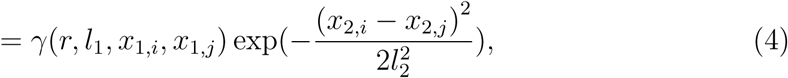

with 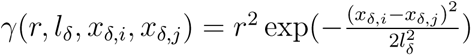 for *δ* = 1, 2. Let us look at how the interpretation of lengthscales *l*_1_ and *l*_2_ can be differently from what we see with 1-dimensional inputs in Equation 2. Specifically, consider reducing *l*_1_ while keeping all other parameters constant:

- **From the perspective of the 1***st* **dimension (Equation 3):** The term *γ*(*r, l*_2_*, x*_2_*_,i_, x*_2_*_,j_*) acts as an unaffected scale factor. In this case, reducing *l*_1_, similar to what is observed in Figure 12 right, results in “more wiggly” curves. A smaller *l*_1_ increases the influence of nearby points on the curve, making their impact more pronounced.
- **From the perspective of the 2***nd* **dimension (Equation 4):** The term *γ*(*r, l*_1_*, x*_1_*_,i_, x*_1_*_,j_*) becomes an affected scale factor. Reducing *l*_1_ lowers this factor due to the decrease in 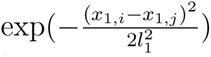. As shown in Figure 12 left, this reduction decreases the covariance between points, regardless of their distances.
- It is also important to note that these perturbations of lengthscales would not affect the variance of the points themselves (i.e. *k_Gau_*(***x_i_***, ***x_i_***)), as scale factor *r*^2^ does not really change.

For a visual representation of the explanations discussed above, please refer to the example provided in Appendix B.2.1.

The exploration in this section demonstrates the difficulties of conducting perturbation analysis in GP models. Specifically, changes to a single hyperparameter can have complex, non-local effects because of interactions with other hyperparameters. In this case, the standard perturbations, which are typically intended to cause minor adjustments, can sometimes lead to unexpectedly large differences, resulting in models that behave very differently. This variability makes it challenge to generalise findings from the standard perturbation.

Such challenges are particularly relevant in the context of ABC, where the effectiveness of summary statistics depends on capturing model sensitivity to parameter changes. To address this, in Section 3, we propose using Negative Log Posterior Probability (NLPP) scores to redefine model similarities. These scores help by summarizing the overall effects of hyperparameter changes on the posterior distribution, reducing the influence of extreme cases and providing a more balanced view of model sensitivities.

#### B.2.1 An example with visual representation

Suppose we have a vector representing current age, denoted as ***a*** = (*a*_1_*, …, a*_13_) with *a*_1_ *< a*_2_ *< … < a*_13_, and a vector of current logarithmic size, denoted as ***z*** = (*z*_1_*, …, z*_14_) with *z*_1_ *< z*_2_ *< … < z*_14_. To create a list of 2-dimensional inputs, we use the meshgrid function to obtain pairs of ***a*** and ***z***. Specifically, this function produce a set of 182 inputs {***x_i_***}*_i_*_=0_*_,…,_*_181_ = {(*x*_1_*_,i_, x*_2_*_,i_*)^⊤^}*_i_*_=0_*_,…,_*_181_ = {(*z*_1_*, a*_1_)^⊤^, (*z*_1_*, a*_2_)^⊤^*, …,* (*z*_1_*, a*_13_)^⊤^, (*z*_2_*, a*_1_)^⊤^*, …,* (*z*_14_*, a*_12_)^⊤^, (*z*_14_*, a*_13_)^⊤^}.

Note that, the input elements of ***x_i_*** *_i_*_=0_*_,…,_*_181_ are intentionally ordered. This arrangement ensures a periodic variation in their Euclidean distances, and help us easier to virtualise systematic patterns in their covariances.

Let us consider the effect of decreasing *l_z_* (the lengthscale for ***z***) while keeping other parameters unchanged. Specifically, we plug ***x_i_*** *_i_*_=0_*_,…,_*_181_ into the Gaussian kernels used by **M1**, **M2** and their perturbed counterparts, respectively, to compute the covariances between the data points ***x*_0_** and the others, ***x_j_*** *_j_*_=1_*_,…,_*_181_. The results, shown in Figure 13, align with our earlier discussion.

- **From the perspective for the dimension of age:** reducing *l_z_* decreases the covariance between *x*_0_ and the other points, regardless of their distance. This effect is evident in the pairs of waves in Figure 13 left, where one such pair is highlighted.
- **From the aspect for the dimension of** *z*: as shown by the peaks of the waves in Figure 13 right, when ***a*** (age) is fixed, points {***x_j_***}*_j_*_=1_*_,…,_*_181_ with *x*^2^ that are far from *x*^2^ become less correlated with ***x*_0_**. In this case, points closer to *x*^2^ thus have a relatively higher impact on it.

From Figure 13, we can see that the perturbation affects both models in a similar way. However, the differences in hyperparameter values between **M1** and **M2** amplify or diminish the impact to varying degrees. This highlights that, in GP models, the effect of perturbing a single hyperparameter depends heavily on the values of the other hyperparameters. Furthermore, from Figure 13, it seems the elasticity analysis (proportional perturbation) is more suitable for GP models, but, again, this still depends on the values of other hyperparameters.

**Figure 13:**
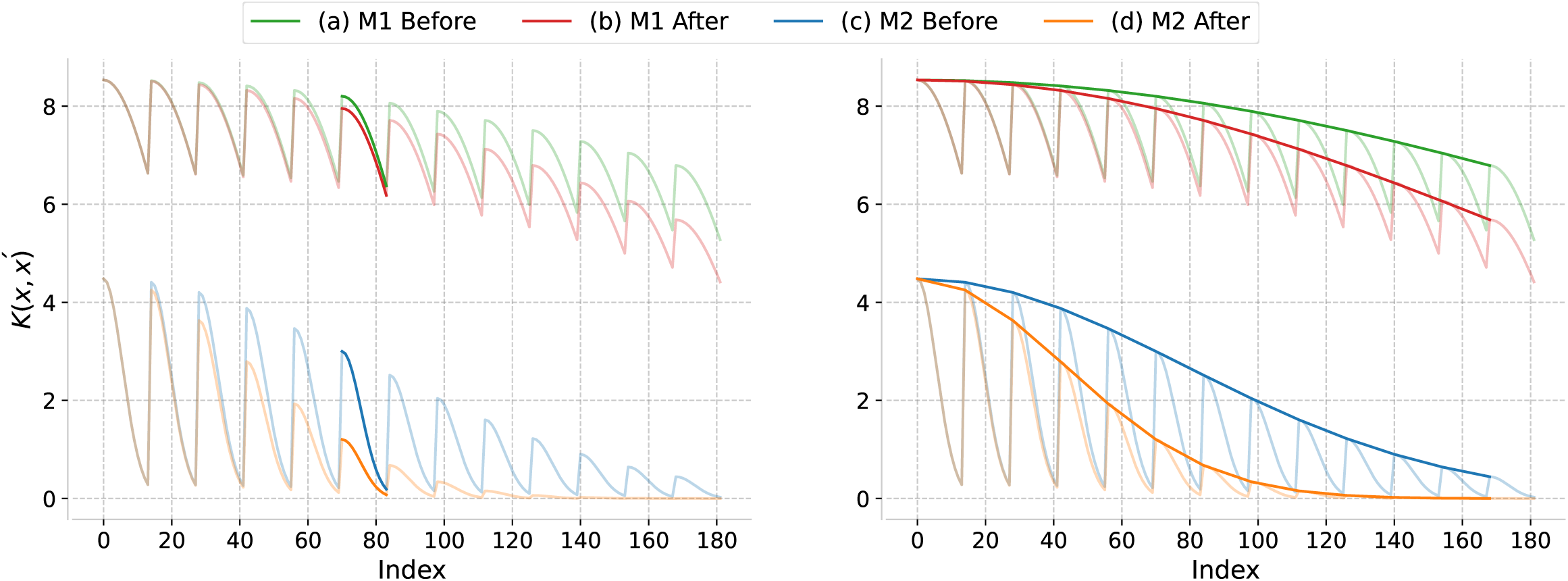
The GP kernel values for data points (***x*_0_**, ***x_j_***) for *j* = 1*, …,* 181. The calculations were performed using Gaussian kernel with four different sets of hyperparameters, which were used in the models that generated the prediction surface depicted in Figure 10. The green and red lines represent the kernel values computed before and after perturbation, respectively, for model **M1**, while the blue and orange lines represent those for model **M2**. The set of values ***x_i_*** *_i_*_=0_*_,…,_*_181_ was obtained through the meshgrid function, resulting in periodic patterns in the plots. The two plots are essentially the same, except that different regions have been highlighted. Notice that, this figure only illustrates how perturbations influence the GP covariance kernels. These changes in covariance also impact the predictive distributions. However, the prediction surface’s response depends on the specific dataset. As a non-parametric method, GP models rely on the positions of the data points to evaluate these effects. Using the conditional density formula for a multivariate normal (MVN) distribution, one may calculate how covariance changes influence predictions in GP regression models.

#### B.2.2 Effects of proportional perturbations

**Table 6:** The values of the hyper-parameters of two GP regression models in Figure 14 and Figure 15 (growth model for breeding individuals). Models producing Plot (b) and (d) were produced by perturbing the lengthscale for the log size with the same proportional amount of noise, −25%, on (a) and (c) respectively.

| Plot | Kernel | NLL | $n_s$ | Scale $r^2$ | ARD | |
| --- | --- | --- | --- | --- | --- | --- |
|  |  |  |  |  | Size | Age |
| (a) | Gau | 54.02 | 0.3243 | 8.533 | $l_1 = 7.098$ | $l_2 = 18.31$ |
| (b) | Gau | 53.94 | 0.3243 | 8.533 | $l_1 = 5.323$ | $l_2 = 18.31$ |
| (c) | Gau | 54.52 | 0.2952 | 4.480 | $l_1 = 2.233$ | $l_2 = 5.531$ |
| (d) | Gau | 55.20 | 0.2952 | 4.480 | $l_1 = 1.675$ | $l_2 = 5.531$ |

**Figure 14:**
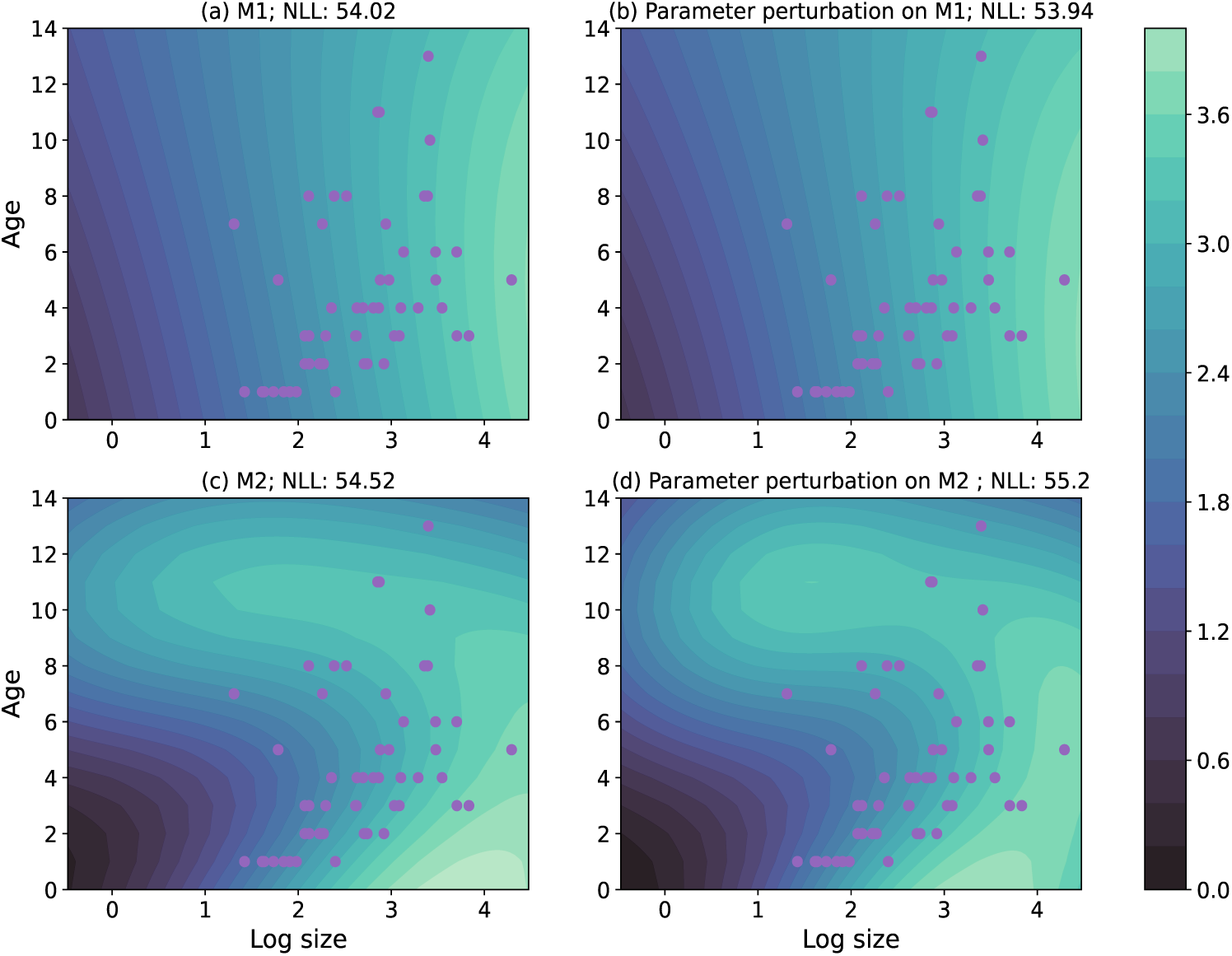
Given the current age (y-axis) logged size (x-axis), the contour plots represent the predicted mean for logged size in the next year. Predictions were made through GP models with hyper-parameters shown in Table 6. Plot (b) and (d) were produced by perturbing the same hyper-parameter with the same **proportional** amount of noise on (a) and (c) respectively. Purple dots represent for the values of covariance for the training data points.

**Figure 15:**
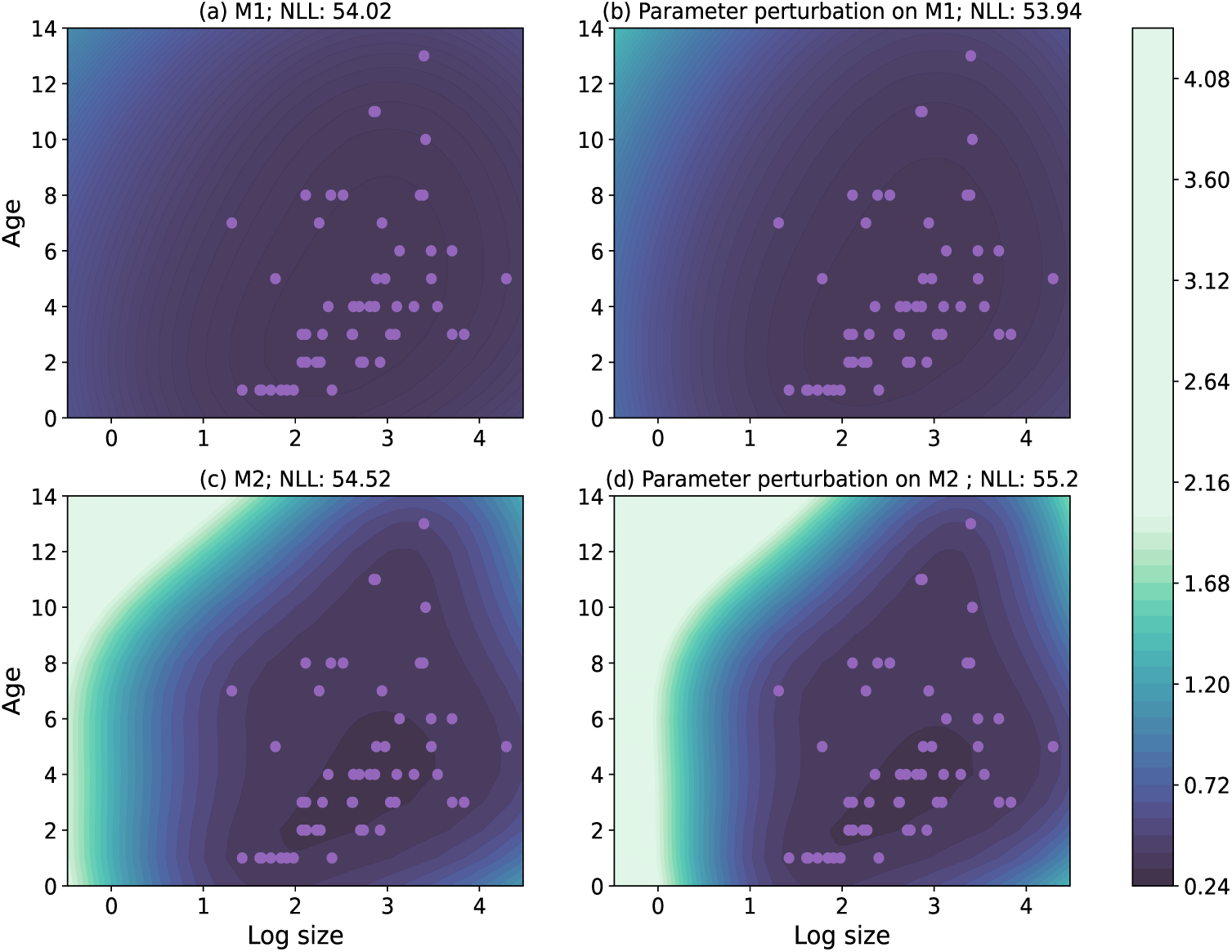
Contour plots for the predicted variance corresponding to the predicted mean shown in Figure 14.

## C The simulation case study

### C.1 Implementation Details

We initiated our experiment by applying GP models and GLMs within a frequentist framework to fit the real-world population data (aggregated from the three years from 2008 to 2011). In this simulation case study, we ignored the ‘age’ dependent and the environmental stochasticity. Specifically, we defined the reproduction kernel as *F* (*z*′|*z*) = *p_b_*(*z*)*n_f_* (*z*)*p_est_r_size_*(*z*′) and the survival-growth kernel as *P* (*z*′|*z*) = *s*(*z*)*p_b_*(*z*)*G_r_*(*z*′|*z*) + *s*(*z*)(1 − *p_b_*(*z*))*G_n_*(*z*′|*z*) (with *z* represents size). These formulations, along with their detailed explanations, were presented in Section 2.2.

To discretize IPM kernels, we employed 50 midpoints to describe the distribution of individual traits, following the default settings in the R package IPMpack (Metcalf et al. (2013)). The lower and upper limits were fixed at 0 and 5 to encompass and slightly exceed the observed log-transformed sizes of the plants (Rees et al. (2014)). To address eviction - where individuals with predicted growth probabilities exceed the limits of the IPM matrices are “accidentally” eliminated - we implemented a ‘constant’ style correction (Williams et al. (2012); Metcalf et al. (2013)).

**Table 7:** The list of the Selected Summary statistics employed when applying ABC-PMC to ABC GP for datasets *D_glm_*and *D_gp_*.

| Selected Summary statistics |  |
| --- | --- |
| $D_{glm}$ | (1) SSD for the total number of breeders in size groups;<br>(2) SSD for the average number of flowering stalks in size groups;<br>(3) the $\chi^2$ distance in total number of individuals in sizeNext groups;<br>(4) the Bhattacharyya distance in non-breeders’ sizeNext distribution;<br>(5) SSD for the total number of survivors in size groups. |
| $D_{gp}$ | (1) the $\chi^2$ distance for the total number of breeders in size groups;<br>(2) SSD for the average number of flowering stalks in size groups;<br>(3) the EMD in breeders’ sizeNext distribution;<br>(4) the Hilbert projective metrice in breeders’ and non-breeders’ survival distribution;<br>(5) SSD for the total number of survivors in size groups. |

For each of GP models in this experiment, the mean function was set to zero (as in common practices), and the most widely used kernel function, the Squared Exponential, was applied (Williams & Rasmussen (2006)). Bayesian individual vital rate model fitting was conducted using non-informative priors and Hamiltonian Monte Carlo methods (Hensman et al. (2015)). Frequentist individual model fitting for GP regressions can be computed directly, while that for GP models with non-Gaussian likelihood, Variational inference (Opper & Archambeau (2009)) was employed. All the standard GP and GLM model fitting tasks were implemented using the GPflow, statsmodels and Bambi package in the Python programming language (Van Rossum & Drake Jr (1995); Seabold & Perktold (2010); Matthews et al. (2017); Capretto et al. (2022)). Workflow 1 was executed for each vital rate model, ensuring that there was at least one summary statistic that performed well for each vital rate. We retained only the most robust summary statistics after implementing Workflow 1, resulting in five different types of summary statistics for each datasets (see Table 7 for details).

**Table 8:** Predictive performance evaluations for the 3 model fitting methods for dataset *D_glm_*.

| Measure | GLM | GP | ABC_GP |
| --- | --- | --- | --- |
| MSE in $\lambda$ | 0.002757 | 0.001899 | 0.000547 |
| Bias in $\lambda$ | 0.04122 | 0.03356 | -0.00033 |
| Mean KL in $\bar{\omega}$ | 0.0104 | 0.0079 | 0.0072 |
| Mean KL in $\bar{\omega}_{rep}$ | 0.0161 | 0.0128 | 0.0119 |
| Mean Cosine similarity in $\gamma$ | 0.9980 | 0.9954 | 0.9953 |

When implementing ABC sampler, for *D_gp_*, {*α_i_*}_0⩽_*_i_*_⩽_*_C_*_−1_ = {0.55, 0.5, 0.45, 0.4, 0.35, 0.3, 0.2, 0.15, 0.1, 0.088; for *D_glm_*, *α_i_* _0⩽_*_i_*_⩽_*_C_*_−1_ = 0.55, 0.5, 0.45, 0.4, 0.35, 0.3, 0.2. For *D_glm_*, the sampler stopped early because no obvious difference was found in the posterior marginal distributions (see Figure 17).

**Table 9:** Predictive performance evaluations for the 3 model fitting methods for dataset *D_gp_*.

| Measure | GLM | GP | ABC_GP |
| --- | --- | --- | --- |
| MSE in $\lambda$ | 0.003129 | 0.001626 | 0.000623 |
| Bias in $\lambda$ | 0.034505 | 0.019512 | 0.009555 |
| Mean KL in $\bar{\omega}$ | 0.0097 | 0.0037 | 0.0018 |
| Mean KL in $\bar{\omega}_{rep}$ | 0.0249 | 0.0140 | 0.0122 |
| Mean Cosine similarity in $\gamma$ | 0.9764 | 0.9947 | 0.9943 |

**Figure 16:**
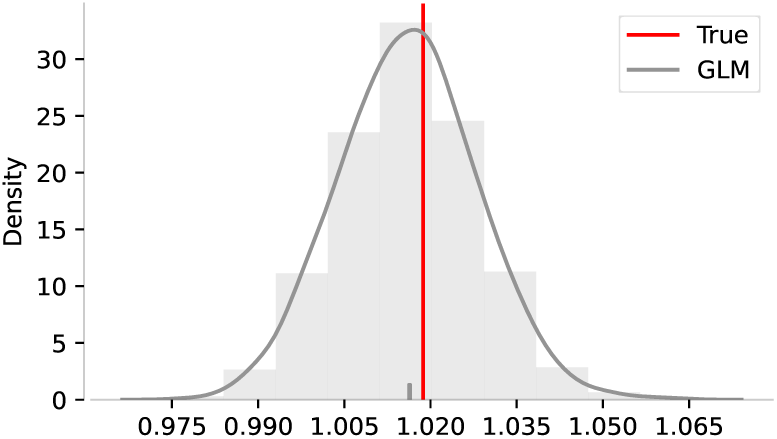
GLM based IPM predictions with a larger sized training dataset, *D_glm_*_2_. Predicted *λ*s, along with their corresponding density histograms (shown as shaded areas), were generated using 5,000 samples draw from the estimated posterior distribution of GLMs. The red line represents the *λ* computed from the true discretized IPM. The pipe (‘|’) in the bottom of each plot marks the corresponding mean.

**Figure 17:**
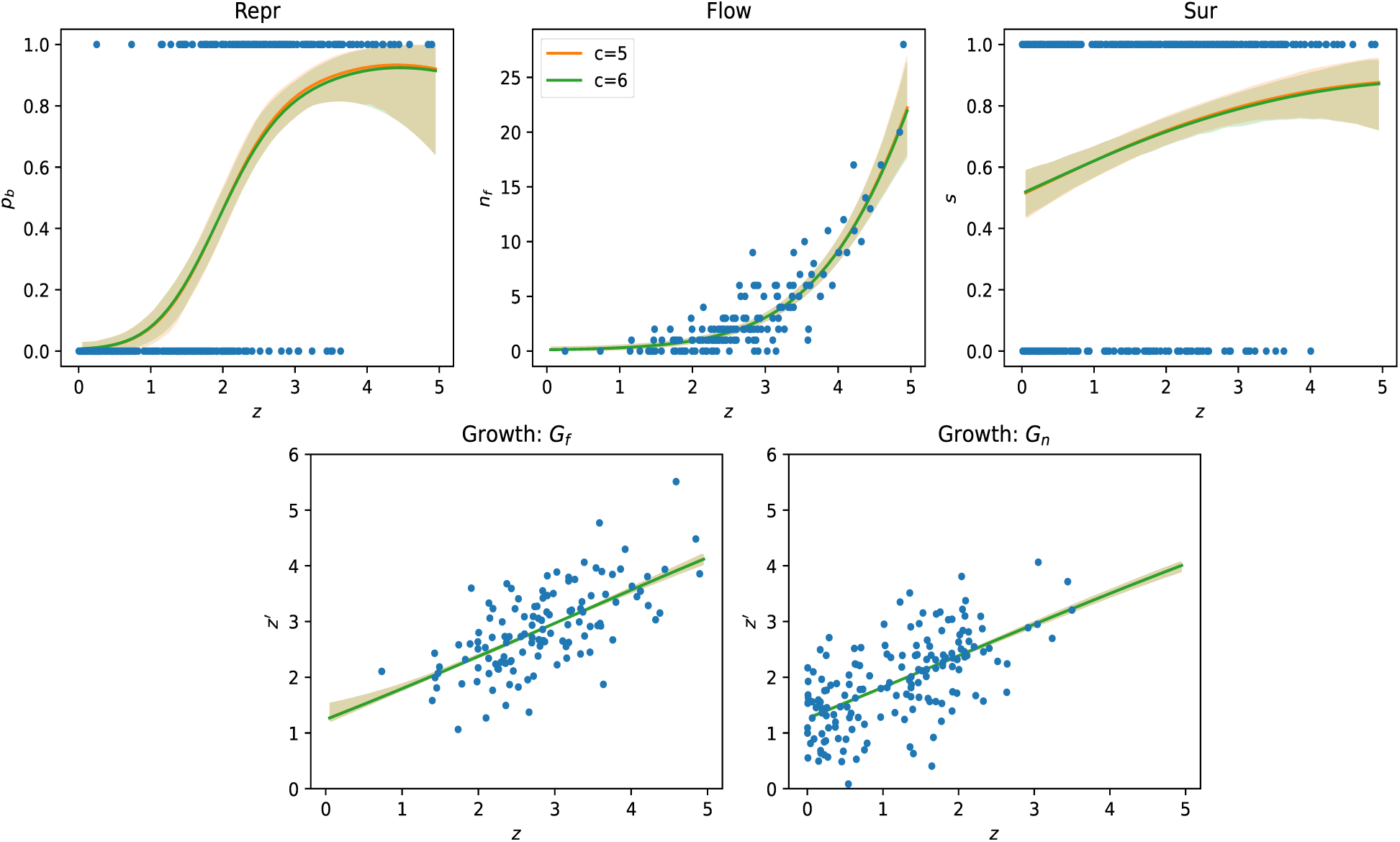
Model fitting results for *D_glm_* at simulation step 5 and 6. The estimated marginal distributions for the five vital rates, complete with their 95% confidence intervals in matching colours and the corresponding scatters data points in blue.

**Figure 18:**
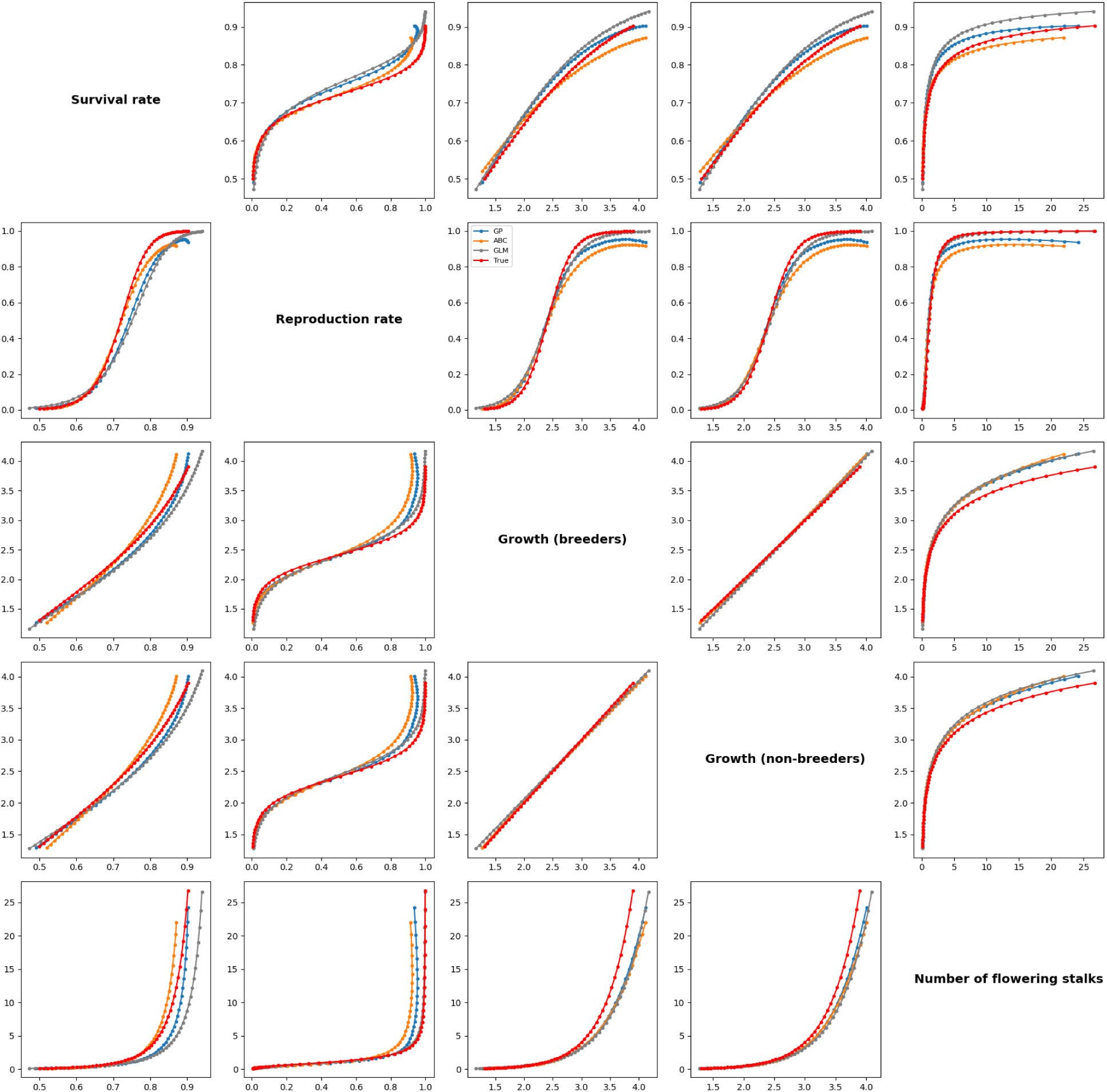
Using the 5,000 samples that generated predicted *λ* in Figure 3 left, we extracted the corresponding vital rate models and evaluated them on 50 mesh points (representing size from 0 to 5). Sample means of vital rates at each mesh point, shown for GLM, GP, ABC-GP, and the true model.

### C.2 Empirical comparison of standard perturbation and NLPP perturbation in GP models

To provide additional context for the use of NLPP-based perturbation in our study, we conducted an experiment to examine how standard perturbation behave in GP models (using the simulated dataset *D_gp_*). Although such perturbations are widely used in ABC methods, it remains unclear whether changes of similar magnitude in parameter space result in comparable model fit among GP models.

If changes in hyperparameters were a reliable inductor for changes in model fit, we would expect a clear pattern — larger perturbations resulting in larger changes in fit, and smaller perturbations yielding similar performance.

#### C.2.1 Experimental setup

For this experiment, we employed a multivariate Gaussian perturbation kernel, which is one of the most commonly used kernels in ABC algorithms (Lintusaari et al. (2017)). The mean vector was constructed by taking the empirical mean of each hyperparameter across the MCMC samples. We choose a diagonal covariance matrix, with each diagonal element estimated as the empirical variance of the corresponding hyperparameter from the MCMC samples of the five vital rate models. (It is worth to notice that, this diagonal structure ignores potential posterior correlations between hyperparameters. While the diagonal simplification may not fully capture the shape of the joint posterior, estimating a reliable full covariance matrix is often challenging in practice, especially in high-dimensional or highly non-linear models. The use of a diagonal approximation is, thereby, very common in ABC applications, for example, Toni et al. (2009); Scranton et al. (2014))

We identified the MCMC samples with the lowest total NLPP across the five vital rate models. This joint configuration, comprising one set of hyperparameters for each rate, was selected as the optimal sample. Using the Gaussian kernel, we generated 5,000 perturbed hyperparameter vectors. For each, we computed:

1. the sum of absolute differences in hyperparameters compared to the optimal sample (across all five models),
2. the sum of absolute differences in NLPP values across the five vital rate models, where each difference is computed by evaluating the perturbed and optimal hyperparameters using their respective GP models on the same data.

While the sum of absolute differences offers a simple and interpretable measure of deviation in parameter space, it does not account for differences in parameter scale or sensitivity. It is used here as a general indicator of perturbation magnitude.

#### C.2.2 Results

Figure 19 Left shows that there is no evidence about linear association between changes in NLPP and the magnitude of hyperparameter perturbation, with a correlation coefficient of −0.0527. Notably, even when the NLPP differences (x-axis) are similar — for example, near 440 — the corresponding hyperparameter differences (y-axis) can vary substantially. This suggests that similar levels of model fit can result from quite different parameter values.

To provide a point of reference, Figure 19 Right shows the distribution of NLPP differences among the original MCMC samples. As expected, these differences are more concentrated and symmetrically distributed, reflecting the level of fit variability under posterior sampling.

**Figure 19:**
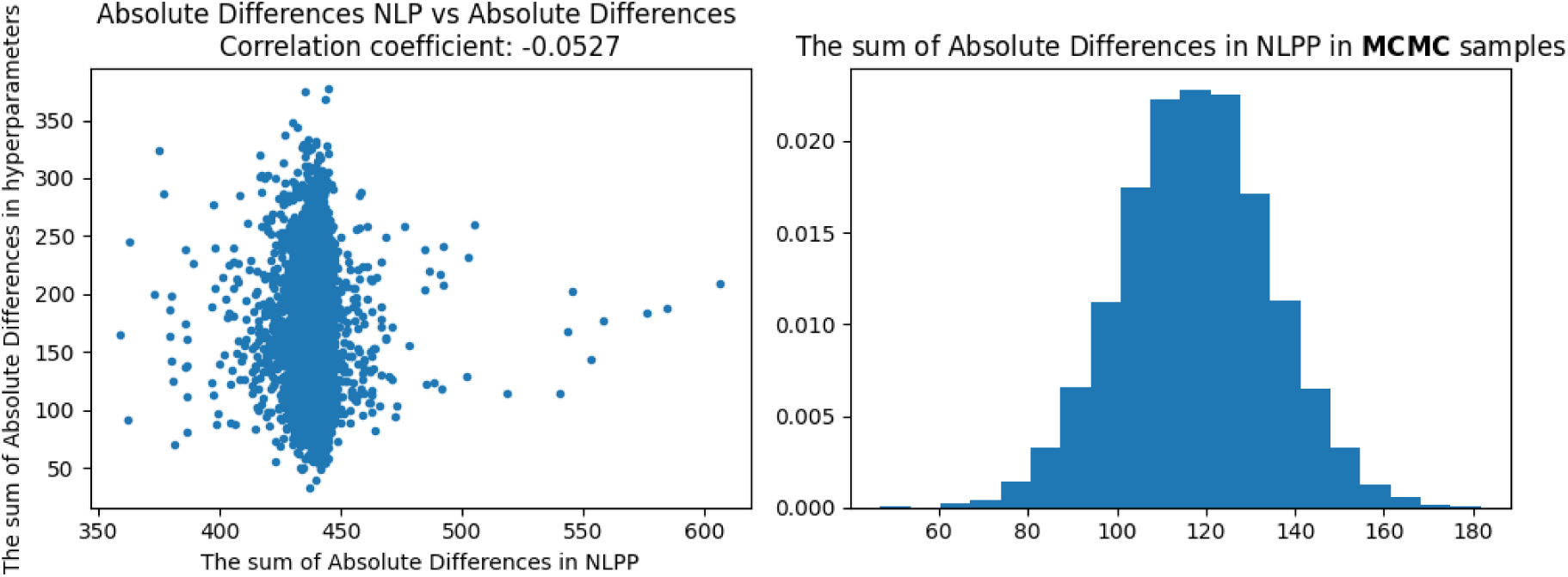
**Left:** the scatter plot showing the sum of absolute differences in hyperparameters and in NLPP values, computed between 5,000 perturbed samples and the optimal sample. **Right:** Density histogram of the sum of absolute differences in NLPP between the optimal sample and randomly drawn 5,000 MCMC samples.

Put these two plot together, what stands out is that none of the 5,000 perturbed samples (Figure 19 Left) reached NLPP values comparable to those from the posterior (Figure 19 Right), even though the perturbation kernel was built using the mean and variance estimated from the same MCMC output and centred around the optimum sample.

In this experiment, we do not observe a clear relationship between the size of hyperparameter perturbation and the change in model fit. While one might expect larger perturbations to lead to larger changes in fit, this is not (consistently) the case here. This observation highlights a practical difficulty in using standard perturbation methods for generating comparably fit in GP models.

As a supplementary check, we also examined the same relationship within the MCMC posterior samples (Figure 20). The correlation between NLPP and hyperparameter differences is similarly low (0.0398), though the range of NLPP values is narrower. This indicate that, in general, differences in hyperparameters are not a reliable indicator of similarity in GP model fit, even within the posterior.

**Figure 20:**
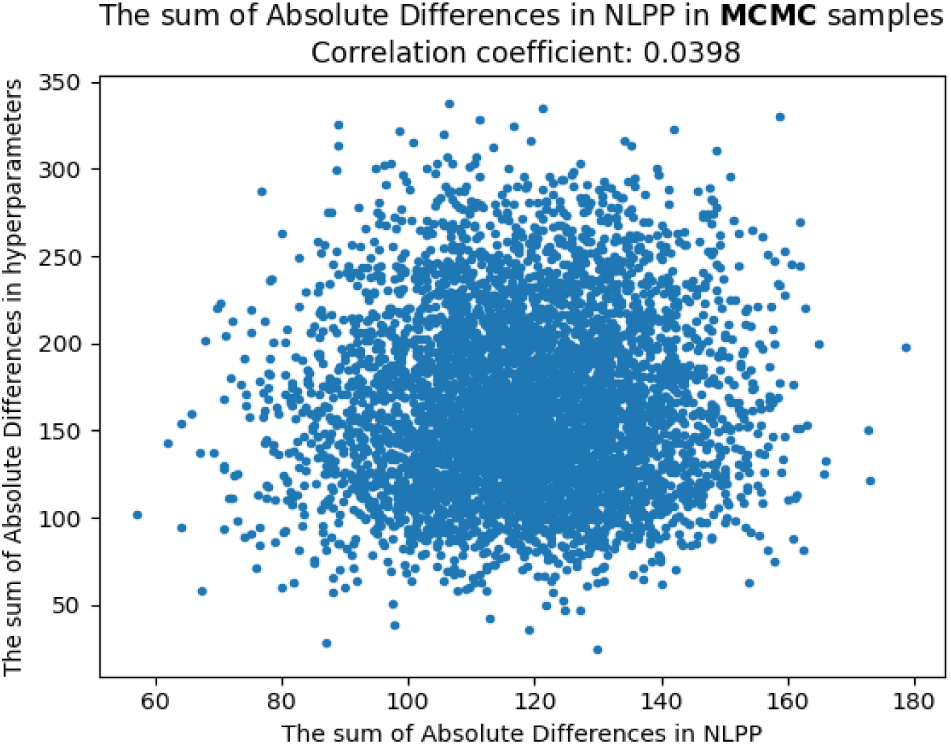
Scatter plot showing the sum of absolute differences in hyperparameters and in NLPP values, computed between between the optimal sample and randomly drawn 5,000 MCMC samples.

## D Real case study

### D.1 Candidate summary statistics

**Table 10:** The list of candidates about the summary statistics and the corresponding distance measures. To calculate the majority of candidate summary statistics under consideration, individuals need to be grouped first according to their size, age, or both. sizeNext and ageNext represent the individuals’ size and age in the next year.

| Summary statistics | Distance measures |
| --- | --- |
| The empirical population distribution in sizeNext, ageNext, sizeNext-ageNext groups | Bhattacharyya distance; Hellinger distance; EMD; K-L distance; Hilbert projective metric |
| Total number of individuals in sizeNext, ageNext, sizeNext-ageNext groups; Total number of flowering stalks in size, age, size-age groups; Total number of breeders in size, age, size-age groups; Total number of survivors in size, age, size-age groups; | $\chi^2$ distance metric; Sum of squared differences |
| Average number of flowering stalks in size, age, size-age groups; Average number of breeders in size, age, size-age groups; Average number of survivors in size, age, size-age groups; | Sum of squared differences |
| breeders' sizeNext and ageNext distribution; non-breeders' sizeNext and ageNext distribution; breeders' and non-breeders' survival distribution | Bhattacharyya distance; Hellinger distance; EMD; K-L distance; Hilbert projective metric |

To calculate the majority of candidate summary statistics under consideration, individuals need to be grouped first according to their size, age, or both. Individuals can be grouped by their current size or age, and can also be categorised by their size or age in the next year (denoted by sizeNext and ageNext respectively). Five size/sizeNext groups are classified using the floor function based on their logarithm size/sizeNext (from 0 to 4). There are also four categories for age/ageNext: age/ageNext = 1 represents newborns; age/ageNext [2, 5] for young individuals; age/ageNext [6, 10] for middle-aged individuals; *>* 10 for white-bearded individuals. With this grouping strategy, for example, the candidate summary statistic ‘the empirical population distribution in sizeNext-ageNext groups’ produces 20 data points. For most of the candidate summary statistics, meanings behind them have already been fully represented by their names. However, we would like to emphasise the following summary statistics that might be slightly confusing:

- **The empirical population distribution in sizeNext, ageNext, sizeNext-ageNext groups:** the counts of individuals in the corresponding groups divided by the total number of individual in the entire population.
- **Average number of flowering stalks/breeders/survivors in size, age, size-age groups:** the counts of flowering stalks/breeders/survivors in the corresponding groups divided by the total number of individuals in the corresponding group.
- **breeders’/non-breeders’ sizeNext and ageNext distribution:** the counts of breeders/non-breeders in the corresponding groups divided by the total number of individual in the entire population.
- **breeders’ and non-breeders’ survival distribution:** individuals are divided into four groups based on whether they were breeders in the past year and whether they survived in the past year.

### D.2 Implementation Details

The model training process will utilise information from the first five years (2003 to 2008), while the data from the last four years will be reserved for model testing. With the dataset’s maximum age being 7 in 2003-2004, we set the absorbing age, *M*, to 8. Individual size, the total number of rosettes, was log-transformed to make it to be less-skewed (in data exploration session; not shown). The yearly population and weather information were altogether plugged into GP models, accounting for the vital rates across the five training years based on 1075 individuals.

Specifically, when *z* represents age and size and sub-kernel *P* and *F* depend on a vector of environmental factors ***ω***, our IPM describes the population of *C. flava* by

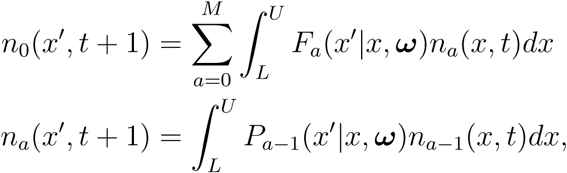

for age *a* = 1, 2*, …, M* 1 and size *x* ∈ [*L, U*]. *P_a_*_−1_(*x*′|*x,* ***ω***) denotes the probability that an individual with age *a* − 1 and size *x* at current time could survive and grow to [*x*′*, x*′ + *dx*] in the next year, with weather conditions ***ω***. Similarly, *F_a_*(*x*′|*x,* ***ω***) describes the expected number of offspring in [*x*′*, x*′ + *dx*] at the next time, per size-*x* individual at current time, with weather conditions ***ω***. *M* is the age class for ‘greybeard’, speaking for individuals with age *M* or older. This requires an addition formula for the model:

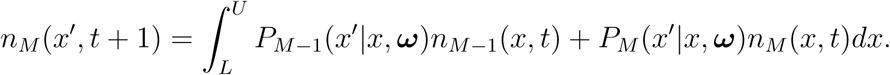

The sequence of life cycle events in the proposed IPMs align with the actual census process conducted in Salguero-Gomez et al. (2012), reflecting realities more properly. Specifically, we defined the reproduction kernel as *F* (*z*′|*z,* ***ω***) = *p_f_* (*z*|***ω***)*n_f_* (*z*|***ω***)*r_est_*_|_***_ω_****r_size_*(*z*′|***ω***) and the survival-growth kernel as *P* (*z*′|*z,* ***ω***) = *s*(*z*|***ω***)*p_f_* (*z*|***ω***)*G_f_* (*z*′|*z,* ***ω***)+*s*(*z*|***ω***)(1 − *p_f_* (*z*|***ω***))*G_nf_* (*z*′|*z,* ***ω***) (with *z* represents size and age). These formulations, along with their detailed explanations, were presented in Section 2.2.

For each of the GP models, the mean function was set to zero, and the most commonly used kernel function, the Squared Exponential, was applied. The Squared Exponential kernel function was employed with the automatic relevance determination setting, allowing it to separately modulate the magnitude of similarity in each input coordinate (e.g. age, size, and various climate factors). Bayesian individual vital rate model fitting was conducted using non-informative priors and Hamiltonian Monte Carlo methods (Hensman et al. (2015)). Frequentist individual model fitting for GP regressions can be computed directly, while that for GP models with non-Gaussian likelihood, Variational inference (Opper & Archambeau (2009)) was employed. All the standard GP model fitting tasks were implemented using the GPflow package in the Python programming language (Van Rossum & Drake Jr (1995); Matthews et al. (2017)).

In this real-world case study, recognising that directly accounting for all environmental factors could introduce great complexity and computational burden, we did not apply Workflow 1 to the entire dataset to select summary statistics. Instead, we focused exclusively on the final year (2007-2008) of the training datasets to minimise the influence of fluctuating environmental conditions across multiple years. The final year was selected because the age information was most complete compared to earlier years. MLEs of the GP models were estimated for the vital rates derived from the 2007-2008 dataset. A simulated dataset, based on the fitted GP models, was then generated and used as the “true” dataset within Workflow 1, allowing for a controlled assessment of model performance. After this, Workflow 1 was executed as usual.

Workflow 1 was executed for each vital rate model, ensuring that there was at least one summary statistic that performed well for each vital rate. We retained only the most robust summary statistics after implementing Workflow 1, resulting in five different types of summary statistics. These are: (1) the sum of squared differences for the total number of breeders in size groups; (2) the sum of squared differences for the average number of flowering stalks in size groups; (3) the Hilbert projective metric in breeders’ sizeNext distribution; (4) the EMD in non-breeders’ sizeNext distribution; and (5) the sum of squared differences for the average number of survivors in size groups. We obtained observed datasets for 5 years, yielding a total of 25 summary statistics. For future use, we denote the distances between a simulated and the observed dataset through the list of selected combinations of summary statistics and distance metrics as 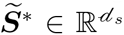 with *d_s_* = 25. When implementing ABC sampler, {*α_i_*}_0⩽_*_i_*_⩽_*_C_*_−1_ = {0.55, 0.5, 0.45, 0.4, 0.35, 0.3, 0.2, 0.15, 0.1, 0.088}; {*N_i_*} with {*N_i_*}_0⩽_*_i_*_⩽_*_C_* = {400000, 240000, 200000, 140000, 100000, 60000, 30000, 20000, 10000, 1600, 200}.

## Notes

### Competing Interest Statement

The authors have declared no competing interest.

### Summary of Updates

We have updated this version to include cross-references to our companion paper.

